# The stoichiometric ‘signature’ of *Rhodopsin*-family G protein-coupled receptors

**DOI:** 10.1101/112466

**Authors:** James H. Felce, Sarah L. Latty, Rachel G. Knox, Susan R. Mattick, Yuan Lui, Steven F. Lee, David Klenerman, Simon J. Davis

## Abstract

Whether *Rhodopsin*-family G protein-coupled receptors (GPCRs) form dimers is highly controversial, with much data both for and against emerging from studies of mostly individual receptors. The types of large-scale comparative studies from which a consensus could eventually emerge have not previously been attempted. Here, we sought to determine the stoichiometric “signatures” of 60 GPCRs expressed by a single human cell-line using orthogonal bioluminescence resonance energy transfer-based and single-molecule microscopy assays. We observed that a relatively small fraction of *Rhodopsin*-family GPCRs behaved as dimers and that these receptors otherwise appeared to be monomeric. Mapped onto the entire family the analysis predicted that fewer than 20% of the ~700 *Rhodopsin*- family receptors form dimers. The clustered distribution of *Rhodopsin-family* dimers, and a striking correlation between receptor stoichiometry and GPCR family-size that we also identified, suggested that evolution has tended to favor the lineage expansion of monomers rather than dimers.

**One Sentence Summary:** Analysis of 71 GPCRs from a single cell reveals the strong tendency of *Rhodopsin*-family receptors to exist as monomers rather than form dimers.

## Introduction

G protein-coupled receptors (GPCRs) are organized into six main families: the *Glutamate, Rhodopsin, Adhesion, Frizzled, Secretin*, and *Taste2* families (*1*). A striking feature of GPCR family structure is the overwhelming dominance of the *Rhodopsin* (class A) family, which comprises >80% of all human GPCRs and a similar fraction of the GPCRs expressed by other vertebrates (*2*). GPCRs all consist of a core of seven transmembrane (TM) α-helices joined by six interhelical loops of variable length. The loops combine with the N- and C-termini forming, respectively, an extracellular region that, together with the TM region creates the ligand-binding site, and a cytoplasmic region that interacts with secondary signaling components, *e.g.* G proteins. The organization of the TM region is strikingly similar across all GPCRs for which structures have been obtained and is stabilized by a conserved network of interactions between topologically equivalent residues (*3*). The most significant structural variation between GPCRs is restricted to the ligand-binding regions, and the parts of the receptors involved in signal-transduction are typically much more highly conserved (*4*), allowing similar conformational changes to accompany receptor activation (5). Several studies of isolated GPCRs (*6–10*) convincingly show that signal transduction can occur on the scale of single, autonomous receptors, consistent with GPCRs forming 1:1 complexes with G proteins (*11*).

Without question the most contentious aspect of GPCR biology concerns their quaternary structures. This is not an insignificant issue as homo- or hetero-oligomer formation offers, *e.g.*, a simple explanation for a wealth of pharmacological data implying that receptors engage in “cross-talk” (although other explanations are possible (*12, 13*)), and new opportunities for pharmacological intervention. Whereas several small families of GPCRs comprise receptors whose large N- and C-terminal domains are known to effect dimerization, *e.g.* the *Glutamate* (class C) receptors (*14*), there is no consensus regarding the “typical” quaternary structure of the largest group of GPCRs, *i.e.* the Rhodopsin-family. It was initially thought that *Rhodopsin-family* GPCRs are generally monomeric, but the more prevalent view now is that these receptors form dimeric and oligomeric complexes with distinct signaling behavior *in vivo* (*15–17*) (the cases for and against oligomerization have been discussed by Bouvier and Hébert and by Lambert and Javitch (*16, 18*)). The first applications of resonance energy transfer (RET)-based assays seemed to precipitate this shift in thinking, but these assays need to be implemented carefully due to difficulties in distinguishing genuine interactions from chance co-localizations, and the interpretation of some early studies is disputed (*12, 19–21*). More recently, single-molecule measurements failed to demonstrate the high levels of constitutive oligomerization expected in transfected and native cells (*22–24*), with one exception (*25*). Equally, lattice contacts in GPCR crystals tend to be reflective of monomeric interactions and, where putative dimers have been observed, the proposed interfaces were not conserved (*26–29*).

Of the RET-based methods, which still afford the highest-resolution (<10 nm) *in situ* assays of receptor organization, bioluminescence resonance energy transfer (BRET) is the most widely used method to study GPCR stoichiometry because it is relatively straightforward and uncomplicated by photobleaching and photoconversion effects confounding Förster RET-based measurements. We previously established three BRET-based assays (types-1 to -3) (*19, 30*), each of which indicates that human β_2_-adrenergic receptor (β_2_AR) and mouse cannabinoid receptor 2 (mCannR2) are monomers. Although these assays were validated using well-characterized monomeric and dimeric protein controls, and correctly identified a known (*Glutamate* family) GPCR dimer, it was not proved by example that native *Rhodopsin*-family dimers were identifiable using these assays. Here, we report a systematic analysis of the stoichiometry of *Rhodopsin*-family GPCRs expressed by a single cell-line using two BRET-based assays and an orthogonal, single-molecule fluorescence-based assay. We observed a very strong tendency for *Rhodopsin*-family to exist as monomers, although we also identified dimers with a high degree of confidence. Combined with an analysis of non *Rhodopsin*-family and root-ancestor GPCRs, our findings suggest a possible explanation for the remarkable asymmetry of GPCR family structure.

## Results

### BRET assay sensitivity

The BRET assays used in this study are described in detail elsewhere (*19, 30*). Briefly, in type-1 BRET experiments (*19*), the ratio of acceptor-to donor-tagged proteins is varied at constant expression and whereas, for monomers, energy transfer efficiency (BRET_*eff*_) is independent of this ratio above a certain threshold, for dimers the relationship is hyperbolic (the principles of the assays are illustrated in Fig. S1). Stoichiometry is indicated by *R*^2^ values for the data fitted to monomer versus dimer models (the principles of our statistical methods are illustrated Figs S2 and S3). In type-3 BRET assays, untagged “competitor” receptors reduce BRET_*eff*_ for dimers but not monomers for a range of expression levels (Fig. S1), with stoichiometry confirmed by the likelihood (*p^diff^*) that the BRET_*eff*_ is affected by the competitor (Fig. S3) (*30*). These assays are complementary insofar as type-1 assays are not prone to false-dimer artifacts but could in principle give false-monomer results in cases of higher-order oligomerization or very weak dimerization, whereas type-3 assays avoid false-monomer results but could produce false dimer signals, *e.g.* when addition of competitor proteins causes the clustering of tagged to be relaxed, reducing effective concentration (Fig. S1). Concordant data obtained with these assays therefore affords confident assignment of receptor stoichiometry. Potential caveats of the assays are discussed in more detail in a Technical Note in the Supplementary Materials.

We undertook a systematic exploration of *Rhodopsin*-family GPCR stoichiometry using both type-1 and -3 BRET assays. By focusing on the set of GPCRs expressed by HEK 293T cells, the host-cell typically used for BRET assays (*15*), we could characterize receptor behavior in its native cellular context. The choice of cell line precluded the use of type-2 assays reliant on observations made at very low expression levels (*19*), because this type of analysis would have been complicated by the presence of untagged, native receptors. The sensitivity of the type-1 and -3 BRET assays was first tested using an inducible system for generating dimers. The monomeric receptor CD86 was fused to the FK506-binding protein (FKBP), allowing the bivalent FKBP ligand AP20187 to induce varying levels of receptor dimerization (Fig. S4). Type-1 and -3 BRET assays detected dimers comprising as few as 10-20% of the total receptor population, as did induced dimerization of β_2_AR (Fig. S4; *R*^2^ and *p^diff^* values are given in Fig. S4C and Dataset S1). For simplicity, we hereafter refer to two classes of receptors: “dimers” (>10-20% dimerization) and “monomers” (0-10% dimerization).

### Two types of *Rhodopsin*-family GPCR behavior

HEK 293T-expressed GPCRs were identified by mining the UniProt database (www.uniprot.org) and comparing the results to gene expression data generated by deep (RNAseq) sequencing of the HEK 293T cell transcriptome. mRNA encoding 65 *Rhodopsin*-family GPCRs was detected in HEK 293T cells (Dataset S2), with assignment of receptors to the *Rhodopsin* family based mostly on published phylogenetic analyses (*1*). These receptors comprised a cross-section of *Rhodopsin*-family GPCRs, with a diverse range of physiological functions and ligands, although some important receptor families were not represented, *e.g.* the dopamine receptors. Transient transfection of GPCRs in the form of GFP fusion proteins in HEK 293T cells gave expression levels of ~100,000/cell (Dataset S1). This included expression on intracellular membranes, consistent with many GPCRs residing mostly in internal membranes until stimulated to traffic to the cell surface (*e.g. 31, 32*). These expression levels were 1-2 orders of magnitude higher than that of native receptors (e.g. *23, 33*), avoiding interference of the assays by homo- and heteromeric interactions with native receptors.

Of the 65 receptors, 60 were judged to have sufficiently good trafficking and expression characteristics to allow BRET analysis (Fig. S5, Table S1). Type-1 and type-3 BRET analysis of 57 of the 60 GPCRs yielded concordant data suggesting that *Rhodopsin*- family GPCRs exist in two stoichiometric states. Representative data sets for monomeric lysophosphatidic (LPA) and dimeric sphingosine-1-phosphate (S1P) receptors are given in Fig. 1A-C. Data for all the receptors are presented in Figure S6. The *R*^2^ and p^diff^ values are plotted in Fig. 1D and absolute values are given in Dataset S1. The results of the analysis, arranged according to *Rhodopsin-family* substructure, are shown schematically in Fig. 1E. For each type-1 assay, total receptor expression was independent of [GFP]/[Rluc] as required by the method (Dataset S1; Figs S6, S7). In this assay, the maximum energy transfer efficiency (BRET_max_) correlated with stoichiometry (*i.e.* higher for dimers than for monomers) and, to a lesser extent with total expression level (Dataset S1; Fig. S8). In the type-3 assay, all dimers gave significantly non-zero projected y-intercepts whereas, for the monomers, the y-intercepts were not significantly non-zero (Dataset S1, Fig. S9). Whilst consistent with our assignments, the *ab initio* interpretation of BRET_max_ and the y-intercepts is not straightforward (see the Technical Note in the Supplementary Materials) and so these metrics were not used to assign stoichiometry to individual receptors.

**Figure 1.**
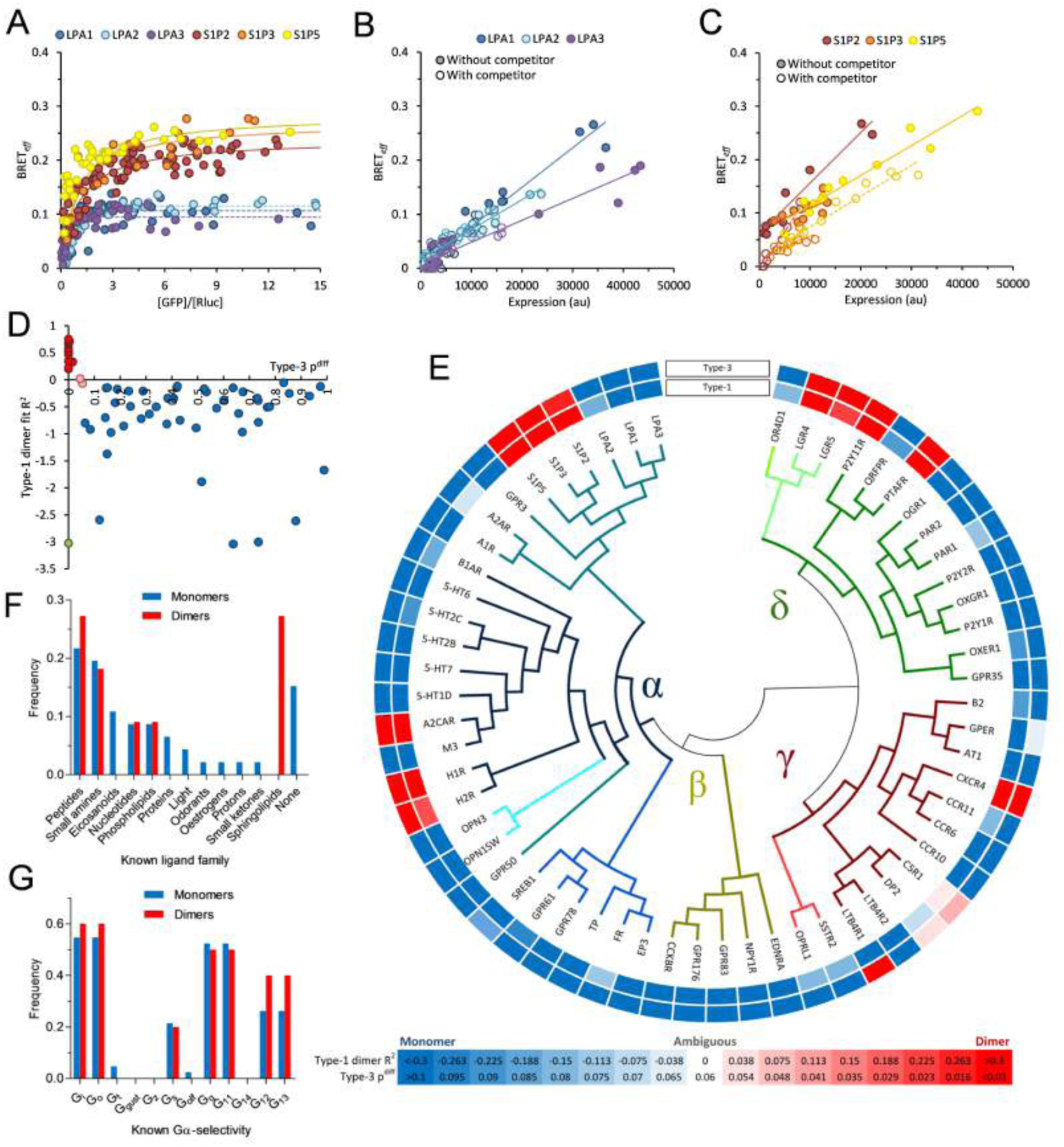
Most *Rhodopsin*-family receptors do not exhibit detectably dimeric behavior in these assays. Representative data for both assays are given for the S1P and LPA receptors, which show contrasting stoichiometry under type-1 (**A**) and type-3 (**B** & **C**) assays. The optimal fit of type-1 data is shown: as a solid line for a dimer model, as a broken line for a monomer model. For type-3 experiments, data collected in both the absence and presence of competitor proteins are shown as filled and empty circles, respectively. For dimers, data fits in the absence of competitor are solid lines whilst those in the presence of competitor are broken lines. For monomers a single fit of all data is shown as a solid line. (**D**) Absolute *R*^2^ (type-1 assay) and p^diff^ (type-3 assay) values for *Rhodopsin*-family GPCRs. Dimeric receptors cluster in the region of high *R*^2^ but low p^diff^. Monomers are shown in blue, dimers in red. C5R1 and DP2 (pink) are ambiguous because they lie near the boundary of monomer vs. dimer outcome under both assays. LTB4R1 (green) is the only receptor to yield conflicting results in the two assays, possibly due to higher-order oligomerization. (**E**) Outcomes for all *Rhodopsin* family receptors investigated summarized as a gradient from blue (monomer) to red (dimer) corresponding to the values given in the key. The inner circle is colored according to type-1 BRET, the outer according to type-3. Receptor relationships are shown as a divergence tree based on previous phylogenetic analysis (*1*). The two receptor populations do not show any clear differences in ligand preference (**F**) or G protein-selectivity (**G**).

Of the 60 receptors investigated, 46 exhibited apparently monomeric behavior in both assays, indicating that they were either wholly monomeric or formed dimers at levels below the sensitivity of the assays. Eleven receptors behaved as dimers in both assays. Two receptors, C5R1 and DP2, yielded data that could not be easily assigned to either form of behavior, possibly due to dimerization at levels near the sensitivity limits of the assays. Only one receptor, LTB4R1, yielded conflicting data in the two assays, perhaps due to higher-order oligomerization, which cannot be unambiguously excluded by the type-1 assay but is readily detected in type-3 assays. Overall, the monomeric and dimeric populations exhibit no obvious differences in either ligand- or G protein-specificity (Fig. 1F,G). Moreover, the stoichiometric assignments do not correlate with C-terminal domain length (Fig. S10A) implying that the cytoplasmic domains did not impose donor-acceptor separation distances greater than the RET-permissive radius (10 nm) leading to the false identification of monomers. Conversely, dimerization is not explained as an artifact of poor trafficking or expression since the fraction of dimers did not correlate with apparent trafficking behavior (Fig. S10B). For the dimers, we are unable to speculate about the strength of dimerization as our analysis generates a binary outcome (*i.e.* fitting more closely to the monomer or dimer model).

### Single-molecule analysis

To further confirm that *Rhodopsin*-family GPCRs do indeed adopt one of two different stoichiometric states, we used single-molecule cross-color coincidence detection (SMCCCD (*34*)). Briefly, candidate GPCRs were transiently expressed in Chinese Hamster Ovary (CHO) K1 cells under the control of a weak promoter to ensure approximately physiological *i.e.* low levels of expression. Unlike HEK 293T cells, CHO K1 cells do not express homologues of any of the receptors studied using SMCCCD (*35*), thereby avoiding interference with the single molecule analysis. Constructs consisting of the receptor fused with a C-terminal HaloTag or a SNAP-tag were co-expressed and then labeled with HaloTag-TMR ligand and SNAP-Cell 505 Star. This allowed individual receptors to be localized in two colors, and the degree of co-localization to be compared to the known monomeric and dimeric controls, CD86 and CD28, respectively. Coincidence values represent the fraction of HaloTag-labeled receptors that localize to within 300 nm of a SNAP-tag-labeled receptor, presented as the mean for all cells analyzed. The principle of the method is summarized in Figure S1 and the Technical Note (see Supplementary Materials).

Receptors exhibiting contrasting behavior in the BRET assays were selected for the SMCCCD analysis: both the S1P receptor 3 (S1P3) and α_2C_-adrenergic receptor (α_2C_AR) behaved as dimers, whereas LPA receptor 1 (LPA1) and β_1_-adrenergic receptor (β_1_AR) exhibited monomeric behavior. Consistent with the BRET analysis, in the single-molecule assay S1P3 and α_2C_AR exhibited above-background levels of cross-color coincidence characteristic of dimers, whereas the LPA1 and β_1_AR receptors exhibited monomer control-levels of cross-color coincidence (Fig. 2 & Table S2). The measured coincidence level was considerably higher for S1P3 than for α_2C_AR, however, which was only slightly higher than that for the monomer control, CD86. It therefore seems likely that, at physiological expression levels, α_2C_AR is a weak dimer).

**Figure 2.**
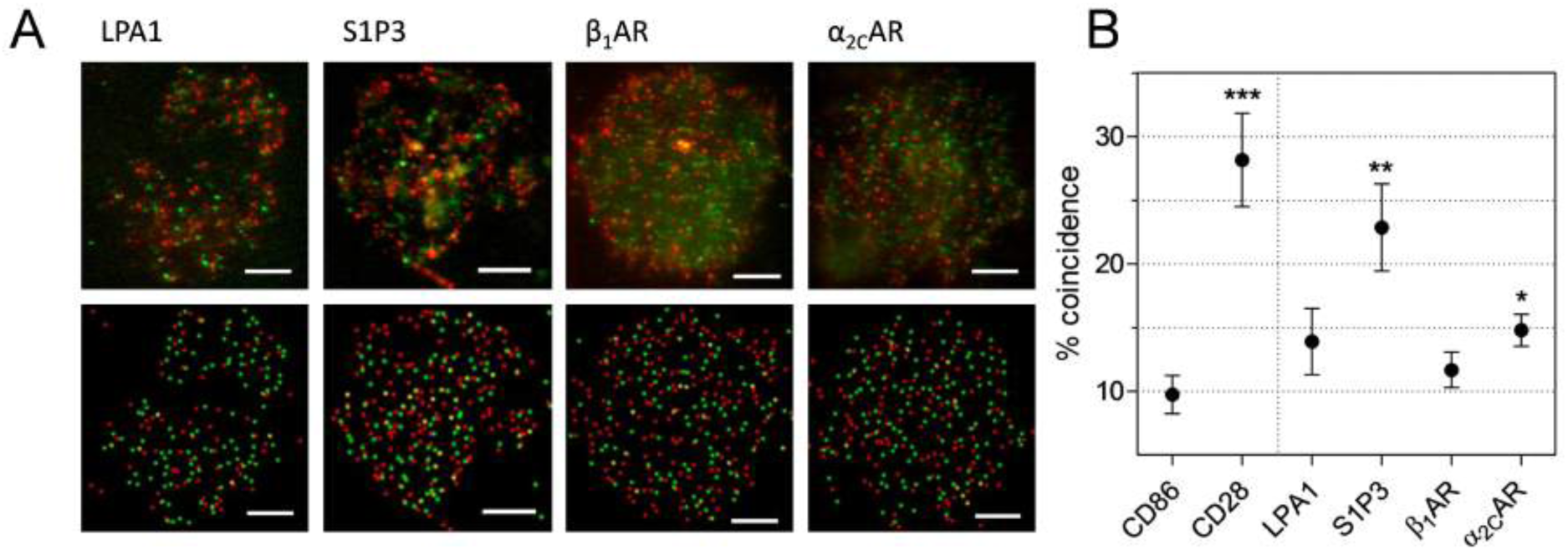
Single-molecule microscopy confirms existence of two stoichiometric states for *Rhodopsin*-family GPCRs. LPA1, S1P3, β_1_AR, and α_2C_AR were expressed as C-terminal SNAP-tag- and HaloTag-fusions to allow two-color labeling. (**A**) Representative actual data (top) and reconstructed spot detection (bottom) of transfected HaloTag (red) and SNAP-tag (green) labeled proteins in CHO K1 cells; scale bars are 5*μ*m. (**B**) S1P3 and α_2C_AR exhibit levels of cross-color coincidence significantly higher than coincidence observed for the strictly monomeric protein CD86, but lower than the covalent dimer CD28, consistent with partially dimeric behavior. Coincidence values for both LPA1 and β_1_AR were not significantly higher than CD86, consistent with monomericity. These observations correlate with the receptors’ reported stoichiometry using BRET. Bars are mean percentage of HaloTag spots within 300 nm of a SNAP-tag spot ± SE. *p<0.05, **p<0.01, ***p<0.005 (two-tailed *t* test of difference to CD86).

### Dimerization of S1P3 *via* transmembrane helix 4

The contrasting behavior of the otherwise closely related LPA and S1P subgroups allowed coarse mapping of the dimerization interface. In preliminary experiments, three LPA1-S1P3 chimeras were generated with varying contributions from each receptor (Fig. S11A), and these expressed well enough for BRET analysis (Fig. S11B). Type-1 BRET analysis of the chimeras indicated that the presence of the N-terminal domain, transmembrane helix (TM) 1, intracellular loop (IL) 1, and TM3 of S1P3 were not sufficient to induce LPA1 to form dimers, whereas the inclusion also of extracellular loop (EL) 1, TM3, IL2, and TM4 resulted in chimera dimerization (Fig. S11C & Dataset S1). Additional S1P3 domains had no further effect on receptor stoichiometry. This suggested that the sequence motifs mediating interaction in S1P3 are likely predominantly located within EL1, TM3, IL2, and TM4 regions. The contributions of these regions to S1P3 dimerization were further dissected using reciprocal swaps of the individual domains (Fig. 3A). Type-1 and type-3 BRET assays identified TM4 as the principal site of dimerization, since the transfer of LPA1 TM4 to S1P3 partially abrogated S1P3 dimerization (Fig. 3B,C) and transfer of S1P3 TM4 induced LPA1 dimerization (Fig. 3D,E; Dataset S1).

**Figure 3.**
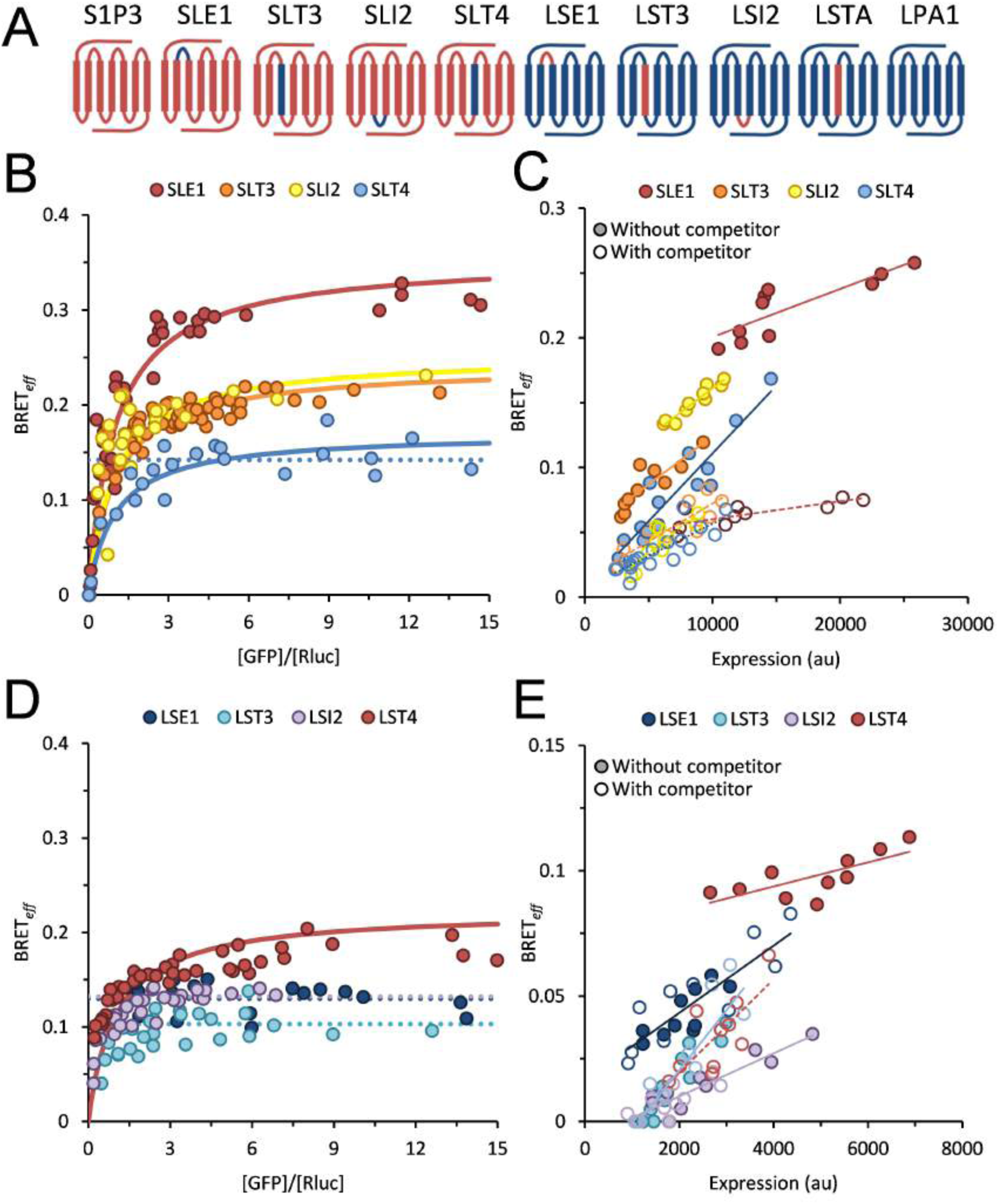
S1P3 dimerization is mediated primarily by interactions involving TM4. (**A**) Schematic representation of the S1P3-LPA1 chimeras studied, indicating their respective composition of S1P3 (red) and LPA1 (blue) sequences. TM helices are arranged 1-7 left to right. Chimeras of S1P3 containing LPA1 domains yielded data consistent with dimerization for all constructs under both type-1 (**B**) and type-3 analysis (**C**). Data collected for SLT4 fit far less closely to a dimer model than the other chimeras, consistent with weaker dimerization. Introduction of S1P3 domains into LPA1 did not induce dimerization of any construct under type-1 (**D**) or type-3 (**E**) analysis with the exception of LST4, indicating that the TM4 domain alone is sufficient to induce receptor dimerization. All data are represented as in Fig. 1.

### Correlation of GPCR family-size and stoichiometry

Our analysis suggests that the majority of *Rhodopsin-family* GPCRs expressed by HEK 293T cells are monomers in contrast to the much smaller set of *Glutamate* family (*14*) receptors, which are constitutive dimers. The phylogenetic distribution of *Rhodopsin*-family dimers that we have identified appears to be non-random, however. Of the eleven dimers, seven are closely related to other dimers (Fig. 1E), forming clusters: *e.g.* the S1P receptors S1P2, S1P3, and S1P5, the histamine receptors H1R and H2R, and the leucine-rich repeat containing receptors LGR4 and LGR5. These observations suggest that the evolutionary appearance of dimers might be rare and episodic. To examine the possibility that stoichiometry might correlate with GPCR family size more generally, we studied the *Frizzled* GPCRs, which appeared contemporaneously with *Rhodopsin*-family GPCRs but comprise only 11 receptors, and *Taste2* GPCRs which emerged and separated from the *Rhodopsin* family just ~300 million years ago but already contain more than 28 members (*36*) exhibiting significant diversification (*37*). Seven *Frizzled* and four *Taste2* HEK 293T-derived GPCRs expressed well enough for BRET analysis (Fig. S5). In type-1 and -3 assays the *Frizzled* receptors all behaved as dimers whereas the *Taste2* receptors exhibited exclusively monomeric behavior (Fig. 4, Dataset S1), suggesting that fast-diverging receptors might generally be monomeric, and that receptors exhibiting less diversification are more often dimers. Chimeric receptors in which the N-terminal, C-terminal, and TM regions of a *Frizzled* (FZD10) and a *Taste2* (TAS2R19) receptor were recombined implicated the N- and C-terminal regions of FZD10 in its dimerization, rather than the TM region (Fig. S4D-F & Dataset S1).

**Figure 4.**
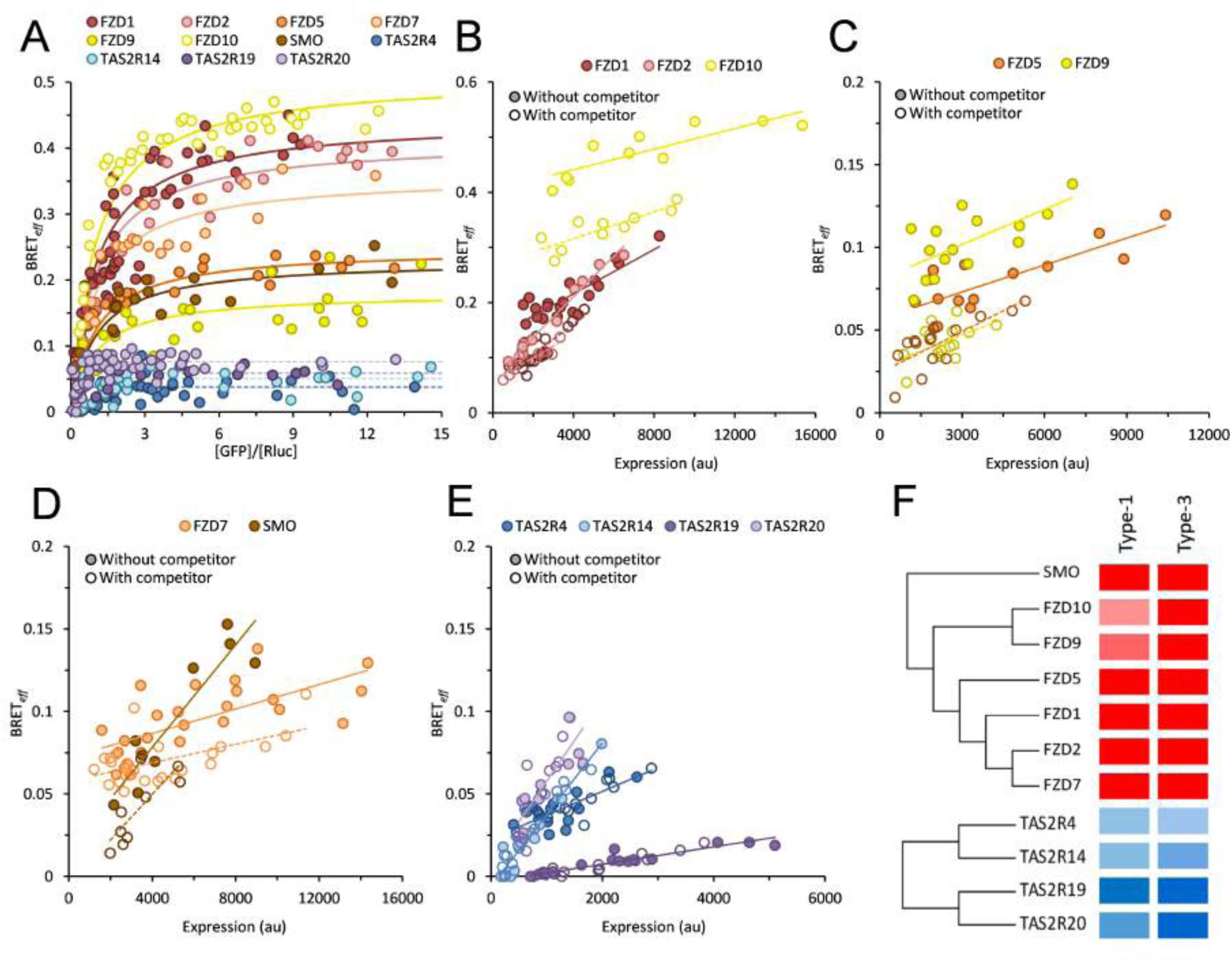
*Fizzled*- and *Taste2*-family GPCRs exhibit contrasting stoichiometry. All seven *Frizzled* receptors studied yielded data consistent with dimeric behavior under both the type-1 (**A**) and type-3 (**B**–**D**) BRET assays. By contrast, the four investigated *Taste2* GPCRs behaved as monomers in type-1 (**A**) and type-3 (**E**) experiments. (**F**) Summarized data according to the color code in Fig. 1. All data are represented as in Fig. 1.

### Stoichiometry of *Rhodopsin* family root-ancestor GPCRs

The *Rhodopsin* family is thought to have emerged ~1.3 billion years ago from the *cAMP* GPCR family (*36*), which was subsequently lost from vertebrates. We examined the stoichiometry of three root ancestor, non-vertebrate *cAMP* family GPCRs in type-1 and type-3 BRET assays. CrlC and CarB are from *Dictyostelium discoideum*, and a receptor we call CLP (*cAMP*-like receptor in Paramecium) is the sole *cAMP*-like GPCR expressed by *Paramecium tetraurelia.* We expect that, because the two species are evolutionarily distant *(D. discoideum* belongs to the amoebozoans, and *P. tetraurelia* in the chromalveolata kingdom), similarities in their behavior will likely reflect the properties of *cAMP* GPCRs generally. All three *cAMP*-family receptors exhibited monomeric behavior in the two BRET assays (Fig. 5, Fig. S5, Dataset S1).

**Figure 5.**
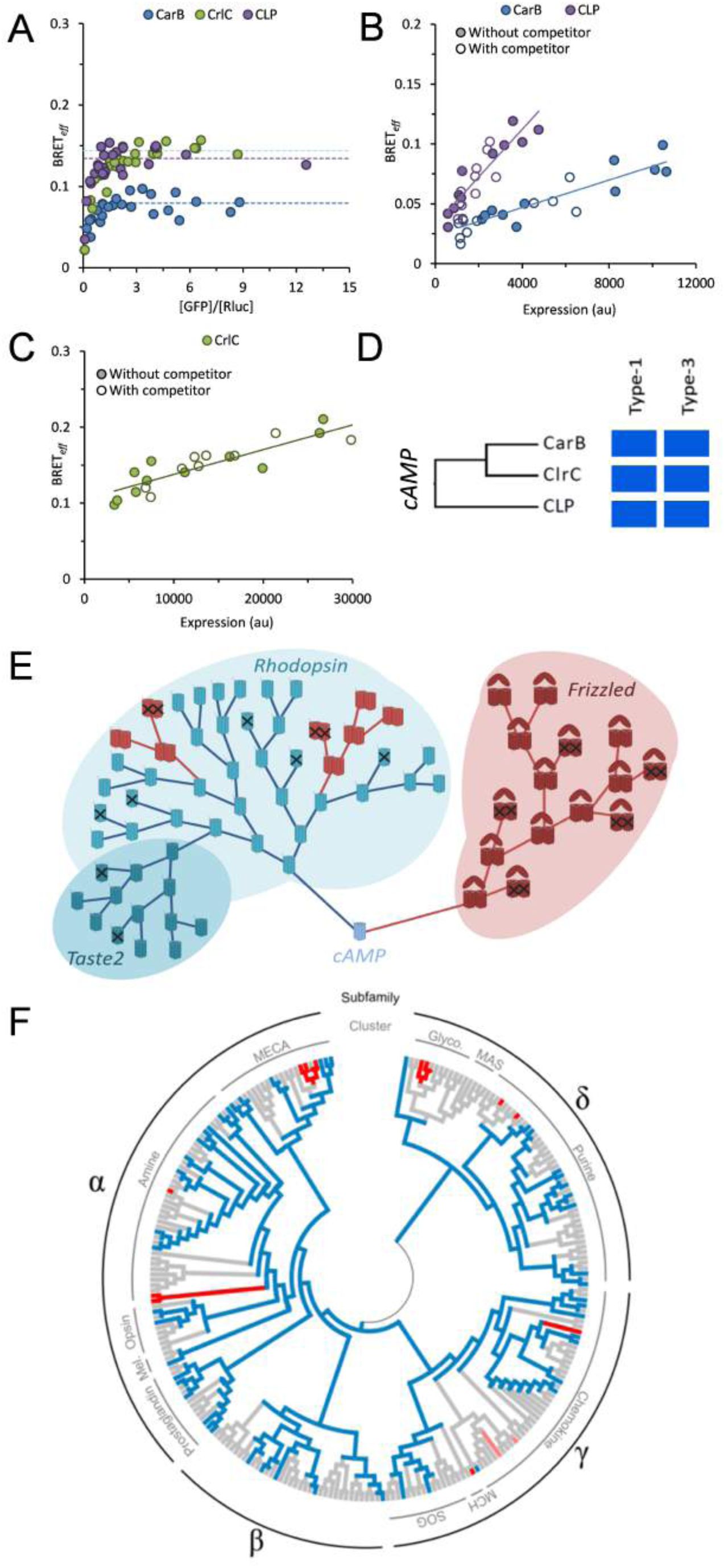
The archetypal stoichiometry of *Rhodopsin-family* GPCRs has been inherited from monomeric ancestors. *cAMP*-family receptors exhibit monomeric behavior under type-1 (**A**) and type-3 (**B** & **C**) BRET assays. All data are represented as in Fig. 1. Data are summarized according to the color code in Fig. 1 in boxed insert (**D**). (**E**) Evolutionary model of GPCR family expansion in which dimers (red) are constrained in their diversification compared to monomers (blue). Receptors that have undergone deleterious mutation are marked with a cross. These receptors are either lost during natural selection, or degrade into pseudogenes over time. Relationships between receptors are represented as a simple lineage tree. (**F**) Gain-of-dimerization events in the *Rhodopsin* family appear to have been episodic. Tree shows all non-olfactory *Rhodopsin* GPCRs along with their subfamily and clusters (*1*). Colored branch end points indicated receptors investigated in this study; red for dimers, blue for monomers, pink for C5R1 and DP2. Grey branches represent receptors not investigated in this study. Nodes are colored in the same way based on their predicted stoichiometry assuming apparent dimers emerged independently during evolution. The distribution of dimers within the *Rhodopsin* family suggests at least seven independent gain-of-dimerization or stabilization-of-dimerization events.

## Discussion

The stoichiometry of Rhodopsin-family GPCRs has been very contentious. Part of the controversy has centered on how best to implement resonance energy transfer measurements in studies of these receptors (*38–40*). Acquired in differently-formatted assays, BRET data was used, for example, to support claims that β_2_AR is an obligate dimer (*41–43*) or that it is constitutively monomeric (*19, 30*). *Rhodopsin-family* monomers and dimers were never identified in any single study, however, leaving open the formal possibility that these assays are incapable of detecting both types of behavior. Here, using two complementary BRET-based assays, we identified multiple examples of *Rhodopsin*-family GPCRs each exhibiting one of two distinct types of behavior, one characteristic of substantive dimers, and the other of monomers. For pairs of monomers and dimers identified in the BRET assays, we observed the same type of behavior in a third, orthogonal single-molecule fluorescence-based assay (SMCCCD). For one of the dimers we tentatively identified the core of the dimerizing interface as TM helix 4 of the receptor. The new data therefore strongly suggest that *Rhodopsin*-family GPCRs are comprised of both monomers and dimers, with the more “typical” behavior being that of a monomer. We have not considered whether the observed oligomerization is functionally significant, however, or whether the relative paucity of *Rhodopsin* family dimers is due to gain-of-dimerization events simply being rare. Although we are also unable to draw direct conclusions regarding heterodimerization or ligand-induced dimerization from our data, we anticipate that the observed resting-state behaviour of the monomers likely reflects a general tendency toward constitutive monomeric behaviour.

Our analysis indicates that as many as 20% of the ~700 *Rhodopsin-family* receptors may form dimers, distributed in discrete clusters across the family (Fig. 5F). We did not attempt to measure the strength of dimerization and it is unclear what fraction of these receptors, if any, dimerize constitutively. S1P3, which, at very low expression levels in the SMCCCD assay produced a coincidence signal indistinguishable from that of our covalent dimer control, CD28, seems to be a good candidate for constitutive receptor homodimerization, however. On the other hand, the BRET measurements were done under conditions of very high expression and it is possible that, due to the effects of mass action, a smaller fraction of *Rhodopsin*-family GPCRs dimerize at lower levels of physiological expression. This type of behavior might be exemplified by α_2C_AR: in the BRET experiments α_2C_AR exhibited clear-cut dimeric behavior, and significantly but only marginally higher levels of co-association than our control monomer, CD86, at the more physiological expression levels of the SMCCCD experiments. For the large set of receptors we sampled using BRET, we are therefore likely to have identified the upper limit of the fraction that form dimers.

Our observations are nevertheless strongly at odds with the notion that *Rhodopsin*-family GPCRs constitutively form dimers. It is noteworthy that of 26 receptors previously reported to be dimers, often in multiple publications (Table S3), 21 behaved as monomers in our assays. The new data are in closer agreement with the growing body of single-molecule microscopy data that has failed to report constitutive dimerization in most cases (*9*, *22*–*24*, *34*). The present data also argue against a general model of allosteric regulation of GPCR homodimers founded, principally, on pharmacological analysis. Instead, cooperative effects observed between receptors might often arise from indirect cross-talk between monomers, as proposed elsewhere (*12*). Our observation that the dimerization of *Frizzled* receptors is apparently mediated by specialized domains outside the TM region offers new support for the notion that the formation of stable dimers requires greater binding energies than can often be provided by the TM region alone, as also suggested elsewhere (*14*). Interestingly, two of the *Rhodopsin*-family GPCR dimers identified here (LGR4 and LGR5) have large extracellular domains similar in structure to those of the *Frizzled* receptors. For the other *Rhodopsin*- family dimers it is more likely that dimerization relies only on TM region contacts, as suggested here for S1P3, implying that these types of receptors might at best interact only transiently.

Several of our observations would be explained by the rare, episodic appearance of *Rhodopsin-family* dimers in the course of vertebrate evolution (Fig. 5E): *(i)* that the dimers comprise a small fraction of GPCRs; *(ii)* that they form small, closely-related phylogenetic clusters; *(iii)* that these receptors do not share ligand- or G protein-selectivity outside the clusters; and *(iv)* that closely related receptors can have different stoichiometries. The infrequency of these clusters seems to mirror a striking correlation between stoichiometry and GPCR family size also suggested by our data. Despite all appearing contemporaneously ~1.3 billion years ago, the dimeric *Frizzled* and *Glutamate* receptor families contain only 11 and 22 members, respectively, whereas there are now >800, predominantly monomeric *Rhodopsin-family* receptors (*36*). Similarly, having split from the *Rhodopsin* family just ~300 million years ago (*36*), it seems remarkable that there are >28 monomeric *Taste2* receptors and only three *Glutamate-family Taste1* dimers (*44, 45*), bearing in mind that both groups of receptors perform the same physiological function.

Why, then, are the *Rhodopsin-family* GPCRs so dominant among extant vertebrates and why are the numbers of putative dimers so low? Assuming that the root ancestor of the *Rhodopsin*-family GPCRs was monomeric, as our initial analysis of the *cAMP* family GPCRs implied, we suggest that at least three forces may have helped shaped *Rhodopsin*-family receptor evolution. First, the functional autonomy of *Rhodopsin*-family monomers (*e.g. 6, 7–9*), allied with an intron-less mode of gene-duplication that would have preserved this functionality, might have favored a classical birth-and-death mechanism of protein evolution (*46*). Second, the cylindrically arranged TM domains, which form deep pockets that bind mostly small ligands, may have allowed very fast functional diversification. Finally, dimerization could have increased the “fitness density” of dimers, constraining the capacity of families of dimers, such as the *Frizzled* and *Glutamate* receptors, to diverge (*47*). This is because, following gene duplication, any “new” receptor would potentially interfere with the function of the “parent” receptor until the capacity for physical interactions was lost, as proposed for other receptor systems (e.g. *47, 48*).

## Materials and methods

### Quantification of HEK 293T transcriptome

The native GPCR expression profile of HEK 293T cells was determined using RNA-Seq. cDNA was generated by reverse transcription of total mRNA harvested from HEK 293T cells, and sequenced using high-throughput RNA-Seq (Illumina). Individual sequences were assigned and quantified using the TopHat and Cufflinks software (Center for Computational Biology, John Hopkins University). Output fragments were then assigned as GPCRs by reference to the Universal Protein Resource (www.uniprot.org).

### BRET vector construction

GPCR genes were cloned into the pGFP^2^ N3 (PerkinElmer), pRluc N3 (PerkinElmer), and pU (described in (*30*)) vectors for BRET. Genes were amplified from cDNA generated from HEK 293T cells by PCR using oligonucleotide primers against the terminal 5’ and 3’ sequences of either the gene open reading frame (ORF) or untranslated regions, or in two stages whereby the 5’ and 3’ halves of the gene ORF were amplified independently and then used as template in the generation of a chimeric full-length product. In some instances, the full length sequence of the gene of interest was synthesised directly by the GeneArt^®^ gene synthesis service (Invitrogen). The *ADRB1* gene was obtained from Robert Lefkowitz, Duke University, in the form of a *FLAG-ADRB1* pcDNA3 vector (*49*) via the Addgene plasmid sharing service (plasmid #14698). This was used as template for full-length PCR amplification as above. The cloning route adopted for each *Rhodopsin*-family GPCR gene is detailed in Dataset S2. Candidate *cAMP*-family receptors were synthesised directly using the GeneArt^®^ gene synthesis service (Invitrogen). FKBP-tagged CD86 and β_2_AR constructs were prepared by the amplification of *FKBP* from the pC_4_-F_V_1E vector (Ariad) using 5’ and 3’ oligonucleotide primers and ligated between *CD86* or *ADRB2* and *GFP*^2^ or *Rluc* in previously described (19) CD86 and β_2_AR N3 BRET vectors.

### Chimeric receptor cloning

Chimeras of the *S1PR3+LPAR1* and *FZD10+TAS2R19* genes were generated using multiple overlapping PCR reactions. Domain boundaries for each gene were identified using the TMHMM v2.0 software from the Centre for Biological Sequence Analysis, Technical University of Denmark. Oligonucleotide primers were designed as complementary to the domain boundaries within the final chimeras, and then used in sequential chimeric PCR reactions.

### Confocal microscopy

Surface expression of GPCRs was assessed via confocal microscopy of their GFP fusion constructs. HEK 293T cells were transfected, 24h after plating onto microscope coverslips, with 1μg pGFP^2^-GPCR DNA per 6×10^5^ cells using GeneJuice^®^ (Novagen) as per the manufacturer’s protocol. Cells were fixed using PBS + 4% para-formaldehyde for 10 minutes. Coverslips were then mounted onto glass slides in the presence of VectaShield mounting medium (Vectorlabs) and imaged using a Zeiss LSM-780 inverted confocal microscope under an oil-immersed 60x objective lens with excitation laser light typically at 488nm. Images were collected at the midpoint of the cell using a 515±15nm emission filter, and minimally manipulated to improve signal-to-noise.

Receptors were assessed qualitatively for cellular distribution and assigned an expression category (Table S1). Protein aggregation was identified by the presence of nonuniform accumulations of GFP fusion protein with fluorescence intensities and *z*-plane distributions greater than that expected of protein retained in internal membranes, as has been demonstrated previously (*50*). Typically, such aggregates would be the brightest objects within the field of view. Genes that demonstrated an expression profile assigned to either category F or G were not progressed to pRluc and pU cloning, nor subsequent BRET analysis.

### Quantitative flow cytometry analysis

Absolute receptor numbers were quantified using flow cytometry analysis of the GFP-tagged receptor variants. HEK 293T cells were transiently transfected with 1*μ*g pGFP^2^-GPCR vector per 6×10^5^ cells using GeneJuice^®^ (Novagen) in the same manner as for the type-1 BRET assay, and incubated for an equivalent length of time as the type-1 BRET assay (*i.e.* 24h for the majority of receptors, 48h for six exceptions). Transfected cells were analysed by flow cytometry for GFP expression. Data were collected for a total of 5×10^4^ cells for each gene in each experiment, and viable single cells were gated for using forward scatter, side scatter, and pulse width. GFP-positive cells were selected using a two-dimensional FL1 vs. FL2 gate to prevent artefacts arising from cellular autofluorescence, and the geometric mean of GFP fluorescence determined for the GFP-positive population.

GFP fluorescence was converted into absolute receptor numbers by reference to a standard curve of GFP fluorescence vs. surface protein expression generated for each experiment using HEK 293T cells expressing a human CD2-GFP fusion protein labeled with phycoerythrin (PE)-conjugated mouse anti-human CD2 antibody (eBioscience 12-0029). Labeling was performed at an antibody concentration of 100*μ*g/ml to ensure saturating, monovalent binding to CD2. PE-GFP compensation was performed using unlabeled cells expressing CD2-GFP as a GFP-only control, and cells expressing CD2-Rluc labeled with PE-antiCD2 as a PE-only control. PE fluorescence on the labeled CD2-GFP cells was then converted to antigen-binding events by reference to calibrated Quantibrite™ (BD Biosciences) PE beads as per the manufacturer’s instructions. Absolute protein numbers were determined in this manner for each receptor in three independent replicate experiments.

### Type-1 and -3 BRET assays

Type-1 and -3 BRET assays were performed on all GPCRs that demonstrated adequate surface expression under confocal microscopy. Type-1 BRET was performed as described previously (*19*) on HEK 293T cells transiently transfected with BRET pairs of the target gene. BRET_*eff*_ and GFP:Rluc ratio were measured 24h post transfection for the majority of receptors, however some GPCRs demonstrated insufficient protein expression to give reliable data after 24h and so were instead measured after 48h. GPCRs measured after 48h were AT_1_, CCR11, GPER, NPY1R, OR4D1, and PAR1. Each receptor was assayed in a minimum of three independent experiments until a broad range of GFP:Rluc ratios had been achieved.

Type-3 BRET assays were performed as described previously (*30*) in HEK 293T cells transiently transfected with BRET pairs of the target gene along with competitor or blank pU expression vector. In all instances, a 2:1 ratio of pU:(pGFP^2^+pRluc) was used to ensure an excess of competitor over labeled proteins, and a 12:1 pGFP^2^: pRluc ratio was used to ensure measurable levels of BRET. In the majority of cases this was achieved using a transfection strategy of 1*μ*g pU, 0.462*μ*g pGFP^2^, and 0.038*μ*g pRluc per well of 6×10^5^ cells, however in cases of low receptor expression these amounts were increased to 2*μ*g pU, 0.924*μ*g pGFP^2^, and 0.076*μ*g pRluc per well. Such an increase in DNA was required for 5-HT_2B_, AT_1_, B_2_, CCR11, EDNRA, GPER, NPY1R, OR4D1, OXER1, and PAR1. Data collected are from a minimum of three independent experiments.

In BRET experiments using FKBP-tagged inducible dimers, cells were incubated for 45 min at room temperature in the presence of varying concentrations of AP20187 in PBS prior to the assay. The concentrations required to achieve varying levels of dimerization were calculated from the relationship between [AP20187] and BRET_*eff*_ for CD86_FKBP_ (Fig. S4A): 0%, 0nM; 10%, 35nM; 20%, 85nM; 30%, 145nM; 40%, 225nM; 50% 335nM; 100%, 5*μ*Μ.

### Statistical analysis of BRET data

Analysis of all BRET data was performed using the Prism5 (Graphpad) software. Type-1 assay data were fitted to models of both dimeric (Equation 1) and monomeric (constant) behavior using the non-linear regression least squares function. The lower and upper range limits of [GFP]/[Rluc] values included in the analysis were 2 and 15, respectively, since 2 is the value at which BRET_*eff*_ becomes independent of acceptor:donor ratio as confirmed with numerous controls (*19*), while 15 is the point at which the dimer model curve has flatted sufficiently to make it indistinguishable from a flat line within the typical error of the experiment. Restricting analysis within these thresholds therefore allows the most sensitive discrimination between monomer and dimer models. Data points between [GFP]/[Rluc] values of 0 and 2 are included in plots for clarity but were not used in either curve fitting or statistical analysis. The coefficient of determination (*R*^2^) for the monomer fit is always zero as calculated BRET_*eff*_ is a constant value. An *R*^2^ value of less than zero for the dimer model indicates that it has a worse goodness-of-fit to the data than the monomer model, whereas an *R*^2^ greater than zero indicates a better goodness-of-fit. Examples of monomer and dimer statistical outcomes are given in Fig. S2.

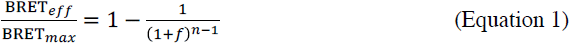

Where:

*f* = acceptor:donor ratio
*n*=stoichiometry
BRET_*max*_ = maximal BRET_*eff*_ achievable in a given system

To confirm protein density did not change with [GFP]/[Rluc], total expression was calculated from absolute luminescence and fluorescence measurements using Equation 2, which expresses protein numbers as arbitrary luminescence units. Total expression *vs.* [GFP]/[Rluc] was plotted and fitted to a linear least-squares regression, then assessed for deviation from non-zero slope using a Fisher F test (*p*<0.05 indicates a significant deviation from zero). *p* values are given in Dataset S1 along with mean % slope as explained in Fig. S7.

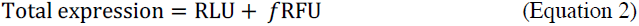

Where:

*f* = acceptor:donor ratio
RLU = relative luminescence units
RFU = relative fluorescence units

All type-3 assay data were fitted using the linear regression least squares function for total expression (from Equation 2) *vs*. BRET_*eff*_, generating separate fits for the data collected in the presence and absence of competitor. Goodness-of-fit was confirmed using the *R*^2^ statistic and found to be high in all cases. *p^diff^* values were determined as the probability that the two datasets were from populations with identical *t* distributions. The larger the *p^diff^* value, the lower the probability of difference between datasets, and hence dimers were defined as those receptors yielding a significant difference between conditions (Fig. S3).

### Vector construction for SMCCD

C-terminally SNAP-tag- and HaloTag-labeled LPA1, S1P3, β_1_AR, and α_2C_AR were generated by subcloning the respective genes from pGFP^2^ into the pHRI-SNAP-tag and pHRI-HaloTag vectors described previously (*34*). In both pHRI vectors, expression is under the control of the ecdysone-dependent minimal promoter such that gene expression is limited to ~2000-4000 per cell. CD86 and CD28 vectors for SMCCD have been described previously (*34*).

### SMCCD analysis

CHO K1 cells were plated and transfected for SMCCD analysis as described previously (*34*). Transfections were performed using the DNA ratios and post-transfection incubation times given in Table S2 to achieve protein densities of 100-1000 HaloTag spots/cell, and SnapTag:HaloTag ratio of 1-6. Sample labeling, fixation, and data collection were performed as described previously (*34*).

## Acknowledgments

This work was supported by the Wellcome Trust and the United Kingdom Medical Research Council. We would like to thank the Royal Society for the University Research Fellowship of S.F.L. (UF120277).

## Author Contributions

J.H.F., S.L.L., D.K., and S.J.D. designed the experiments. J.H.F., S.L.L., S.F.L., D.K., and S.J.D. wrote the manuscript. J.H.F. and R.G.K. cloned, expressed, and collected BRET data for all GPCRs. S.R.M. cloned, expressed, and collected BRET data for most *Frizzled* and *Taste2* GPCRs. J.H.F. analyzed the BRET data. Y.L. performed and analyzed the RNAseq profiling. J.H.F. and S.L.L. performed the microscopy experiments and analyzed the data.

## Supplementary Materials

### Supplementary Discussion

**Figure S1.**
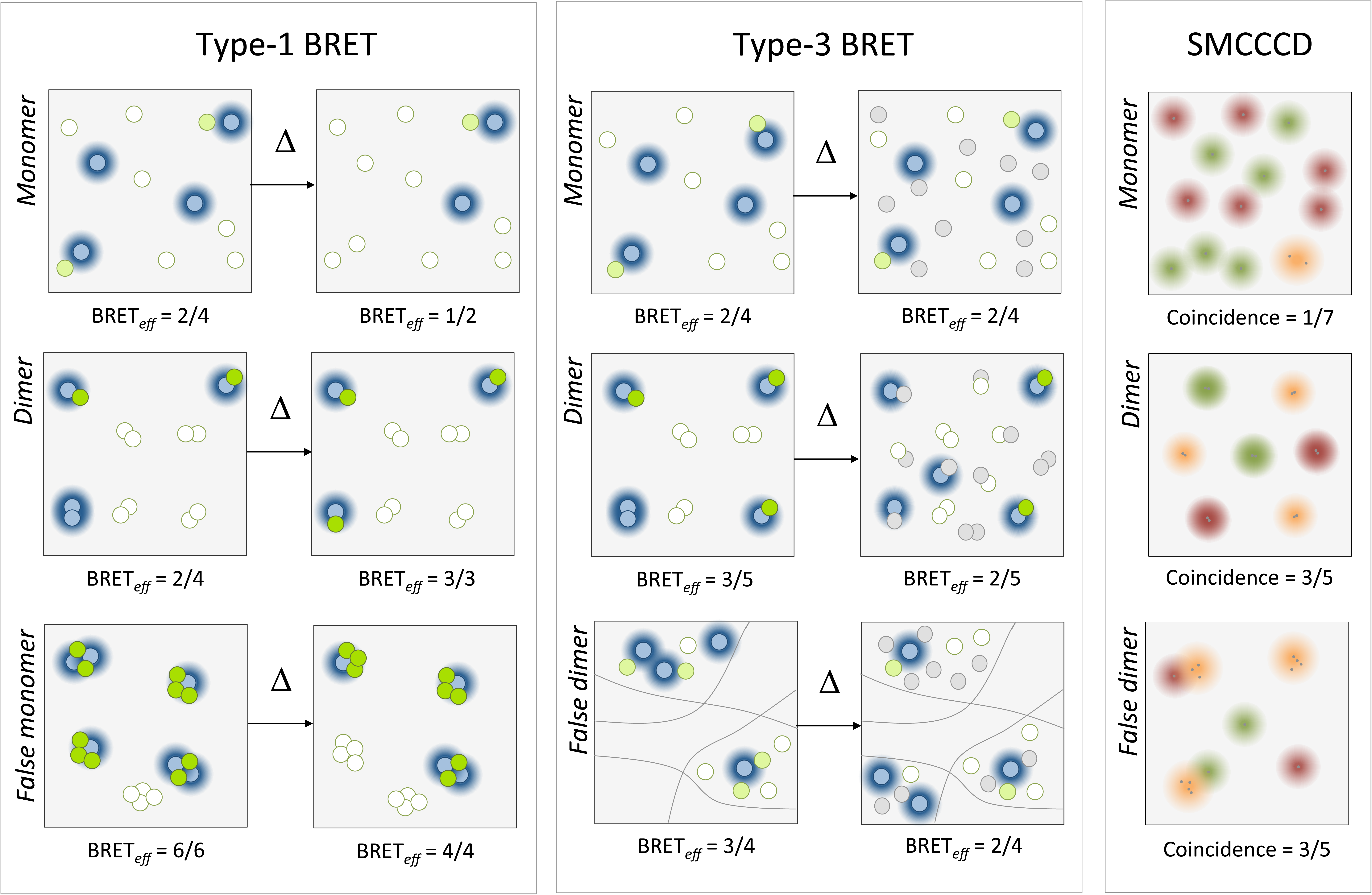
Comparisons of outcomes and limitations of type-1 and -3 BRET assays and SMCCCD. BRET donors are represented as blue circles with halos of the BRET-permissive radius; BRET acceptors as white (non-fluorescing) or green (fluorescing) circles. SMCCCD-imaged fluorophores are shown as individual color (red and green) or combined color (orange) diffraction-radius spots surrounding the tagged protein (gray). In the type-1 BRET assay, monomers exhibit no change in BRET_*eff*_ as acceptor: donor ratio increases because the replacement of donors with acceptors does not impact on the availability of acceptors the remaining donors (left, top). Conversely, as acceptor:donor ratio increases for dimers BRET_*eff*_ will increase as fewer donor-donor pairs form (left, middle). False monomers can be produced in the case of higher-order oligomers as BRET_*eff*_ is also relatively unaffected by increases in acceptor:donor ratio (left, bottom). In the type-3 BRET assay, monomers exhibit no change in BRET_e_f when untagged competitor proteins are introduced into the system (center, top), whereas BRET_*eff*_ for dimers will decrease due to disruption of productive donor-acceptor dimers (center, middle). False dimers can be produced in the case of monomers that undergo clustering within the membrane that becomes more relaxed upon introduction of competitors. This causes a reduction in non-specific BRET_*eff*_ due to reduced effective concentration of tagged proteins (center, bottom). In SMCCCD, tagged proteins are detected as diffraction-limited spots with all proteins within the diffraction limit resolved as a single spot. Proteins are tagged and imaged in two colors, allowing two or more proteins within the diffraction limit (‘coincident’) to be identified. For monomers (right, top), coincident spots are the product only of non-specific colocalization within the diffraction limit. Coincidence is higher for dimers (right, middle) because *bona fide* interaction causes up to 50% (*i.e.* 25% green-green, 25% red-red, 50% green-red) of receptors to colocalize within the diffraction limit. SMCCCD cannot distinguish between genuine dimers and clusters of monomers (right, bottom), which are observed as false dimers.

**Figure S2.**
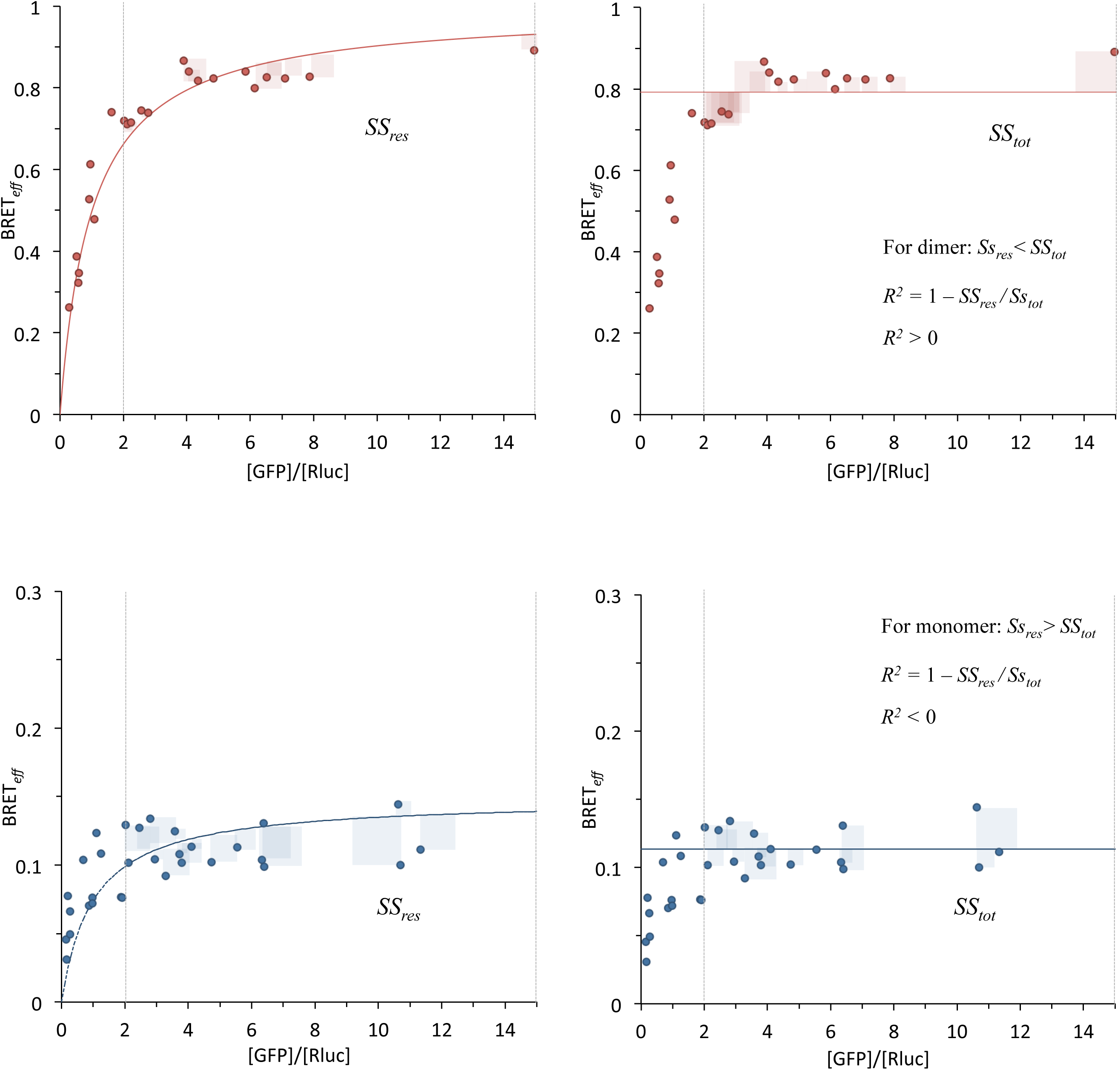
Graphical explanation of type-1 BRET statistical analysis. For all type-1 BRET assays, data were fitted to dimer (left) and monomer (right) models. Fits were generated only for [GFP]/[Rluc] values between 2 and 15. The dimer models fitted data to Equation 1, while the monomer model fitted to a constant BRET_*eff*_ across all [GFP]/[Rluc] values (i.e. BRET_*eff*_ = BRET_*max*_). In both cases non-linear least-squares regression was used to generate optimal fits. To determine which fit better suited the data, the *R*^2^ value was determined. This compares the residual sum of squares for the dimer model fit (*SS_res_*) to that of a flat line (*i.e.* the monomer model; *SS_tot_*)*. R*^2^ is determined as 1 − *SS_res_ / SS_tot_*. For a dimer, *e.g.* CXCR4 (top), *SS_res_* is smaller than *SS_tot_*, so *R*^2^ is positive. For a monomer, *e.g.* CCR6 (bottom), *SS_res_* is larger than *SS_tot_*, so *R*^2^ is negative.

**Figure S3.**
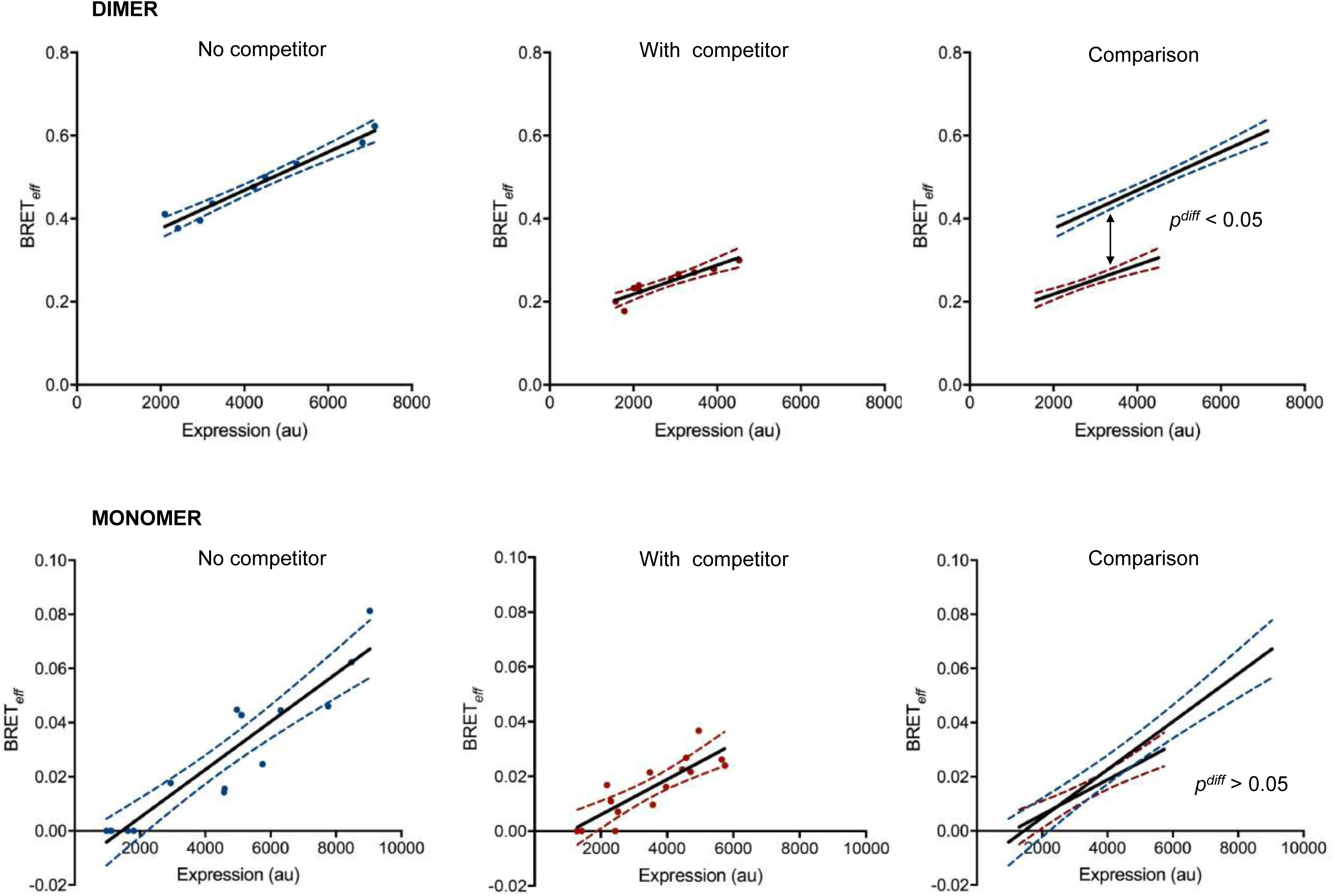
Graphical explanation of type-3 BRET statistical analysis. For all type-3 BRET assays, both datasets (*i.e.* with and without competitor) were fitted to a linear regression least squares model (black line; dotted lines are 95% confidence limits of the fit). All points were used in the generation of the fit. The difference in the elevation of the two fits was then tested using a *t* test to determine the probability that the two fits came from samples with identical *t* distributions. A significant difference in the linear regression models resulted in *p^diff^* value below 0.05, whereas a *p^diff^* over 0.05 indicated no significant difference. The existence of a significant difference between datasets indicated the presence of dimers (e.g. CXCR4; top), whereas its absence suggested monomeric behavior (*e.g.* CCR6; bottom).

**Figure S4.**
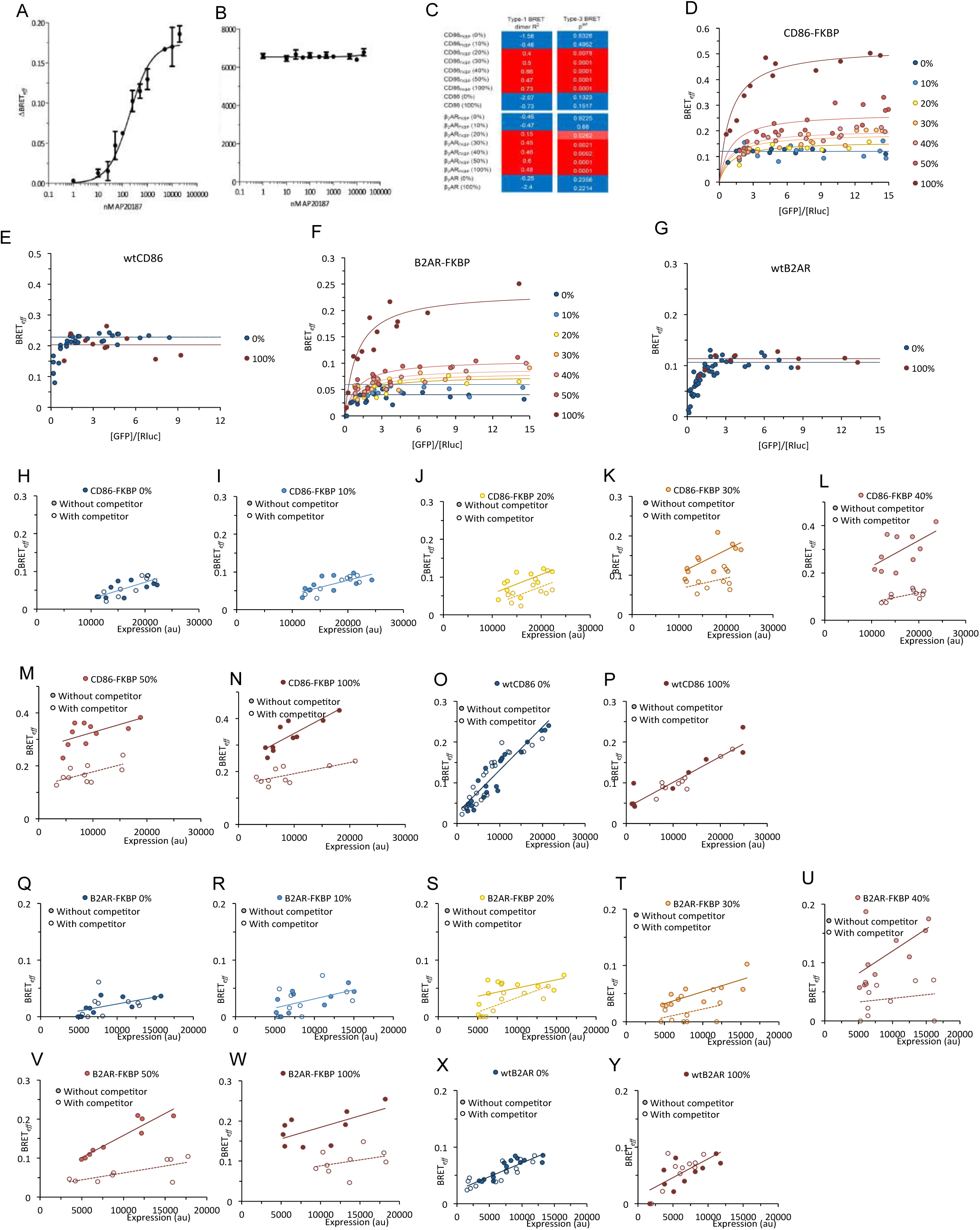
The type-1 and type-3 BRET assays readily detect dimers corresponding to only 20% total protein. (**A**) Change in BRET_*eff*_ between CD86_FKBP_GFP^2^ and CD86_FKBP_**Rluc** in the presence of increasing concentrations of AP20187. AP20187 concentrations required to achieve varying levels of dimerization were derived from these data. (**B**) Total CD86_FKBP_GFP^2^ fluorescence at increasing AP20187 concentrations. Fluorescence is constant across concentrations, indicating AP20187 does not induce internalization and degradation of CD86_FKBP_. The type-1 and type-3 BRET assays can detect dimerization of CD86_FKBP_ and β_2_AR_FKBP_ at AP20187 concentrations sufficient to induce 20% dimerization. (**C**) Summary of outcomes of types-1 and -3 assays on CD86_FKBP_ and β_2_AR_FKBP_ as well as wild-type receptors. Outcomes are colored according to the same code as in Figure 1. (**D** & **F**) CD86_FKBP_ and β_2_AR_FKBP_ demonstrate detectably dimeric behavior in the type-1 assay at AP20187 concentrations sufficient to induce 20% or more dimerization. (**E** & **G**) wtCD86 and wtβ_2_AR exhibit monomeric behavior even at an AP20187 concentration sufficient to induce 100% dimerization of FKBP-tagged equivalents. (**H-N** & **Q-W**) The type-3 BRET assay also detects CD86_FKBP_ and β**2**AR_FKBP_ dimers at 20%. (**O**,**P**,**X**, & **Y**) AP20187 has no effect on the apparent stoichiometry of wtCD86 or wtβ_**2**_AR in the type-3 BRET assay. Data are shown as in Figure 1. Type-1 assay *R*^2^ and type-3 assay *p*^diff^ values for CD86_FKBP_ and β**2**AR_FKBP_ at AP20187 concentrations corresponding to 0, 10, 20, 30, 40, 50, and 100% dimerization are given in Dataset S2.

**Figure S5.**
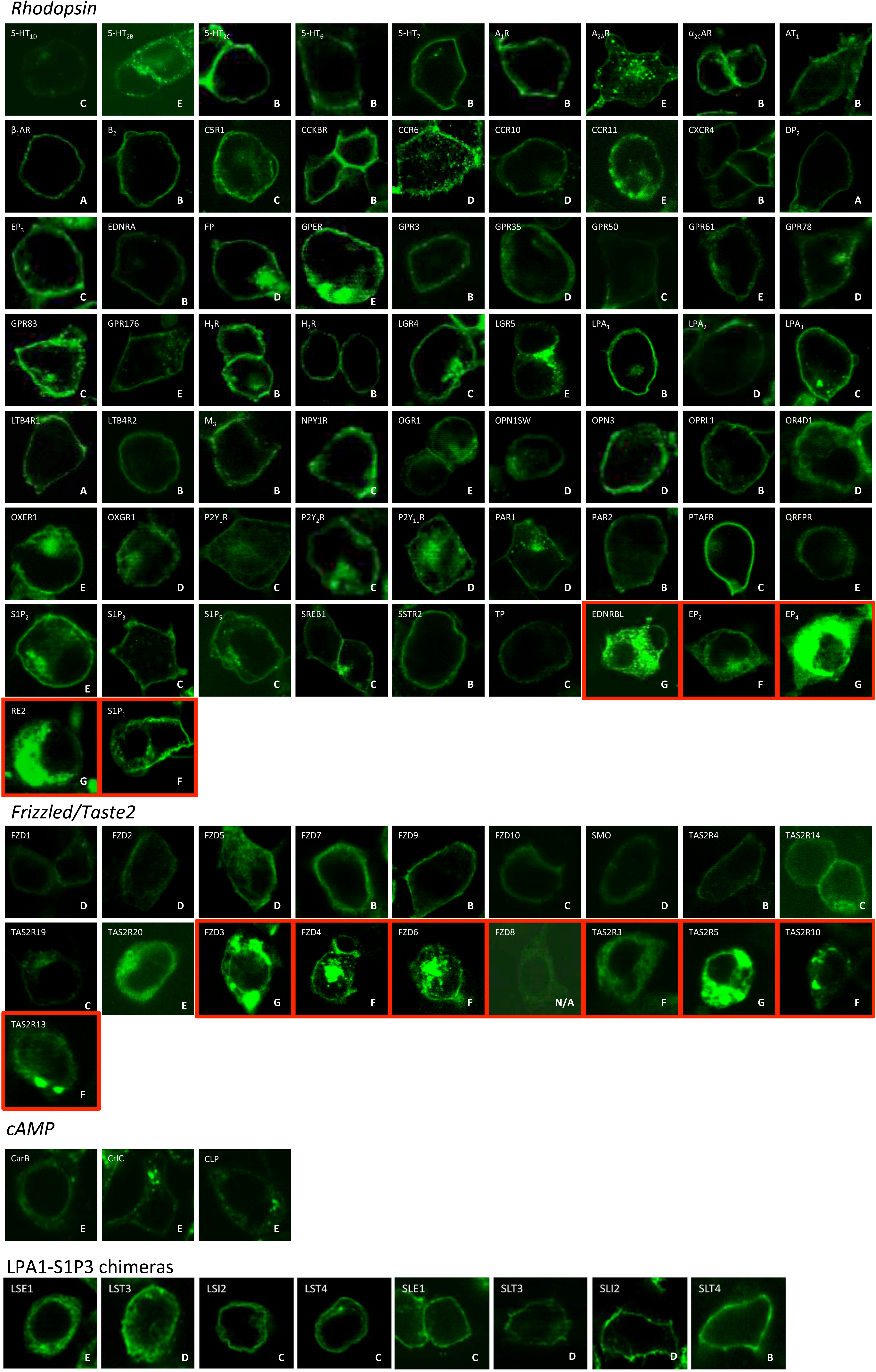
Representative confocal microscopy images of GPCR-GFP constructs expressed in HEK 293T cells. Receptors were placed into categories A-G (Table S1) based on their subcellular localization and degree of observable GFP aggregation. Receptors identified as categories F and G (red borders) were not pursued for BRET. FZD8 expressed too weakly for reliable assessment of localization and so was not pursued for BRET.

**Figure S6.**
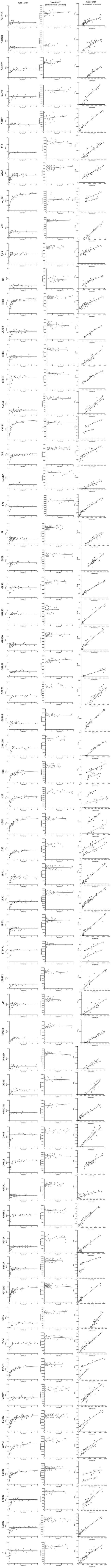
Total datasets for *Rhodopsin*-family GPCRs. Type-1 assay data (left) are shown fitted to either a monomeric (broken line) or dimeric (solid line) model according to which exhibited the best goodness-of-fit. In cases where the outcome was ambiguous both models are shown. Expression *vs.* [GFP]/[Rluc] in type-1 assays are shown fitted to a linear regression (middle). Type-3 assay data (right) are given under conditions of no competitor (gray circles) or with competitor (white circles), with each dataset fitted to linear regression models (solid and broken lines, respectively). All data are summarized in Dataset S1.

**Figure S7.**
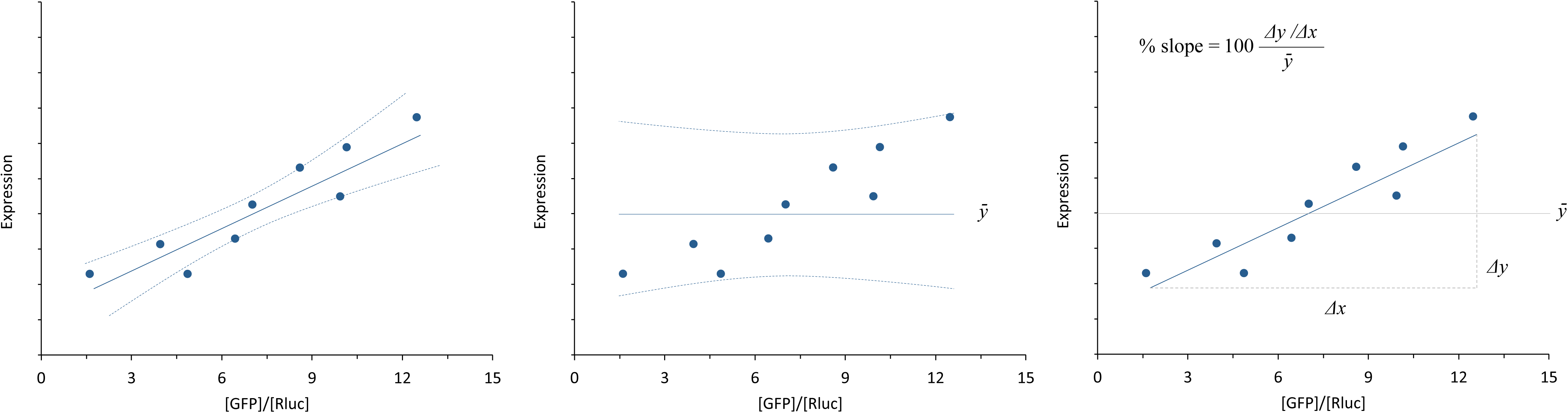
Explanation of *p* and slope metrics for expression *vs.* [GFP]/[Rluc] in type-1 BRET assays. The probability that total protein expression varied systematically with [GFP]/[Rluc] was tested by comparing the goodness-of-fit of a least-squares linear regression fit of the data (left) to that of a zero slope fit around mean expression (middle) using a Fisher test. If the linear regression is significantly non-zero in its slope then the resulting *p* value is <0.05. *P* values for all type-1 BRET assays are provided in Dataset S1, along with mean percentage slope for all samples is also provided. This was calculated as slope exhibited by the linear regression fit expressed as a percentage of the mean expression value for all points (right). *i.e.*, if the percentage slope is 5.00, the total expression would increase 65% across the active range of 2-15 [GFP]/[Rluc].

**Figure S8.**
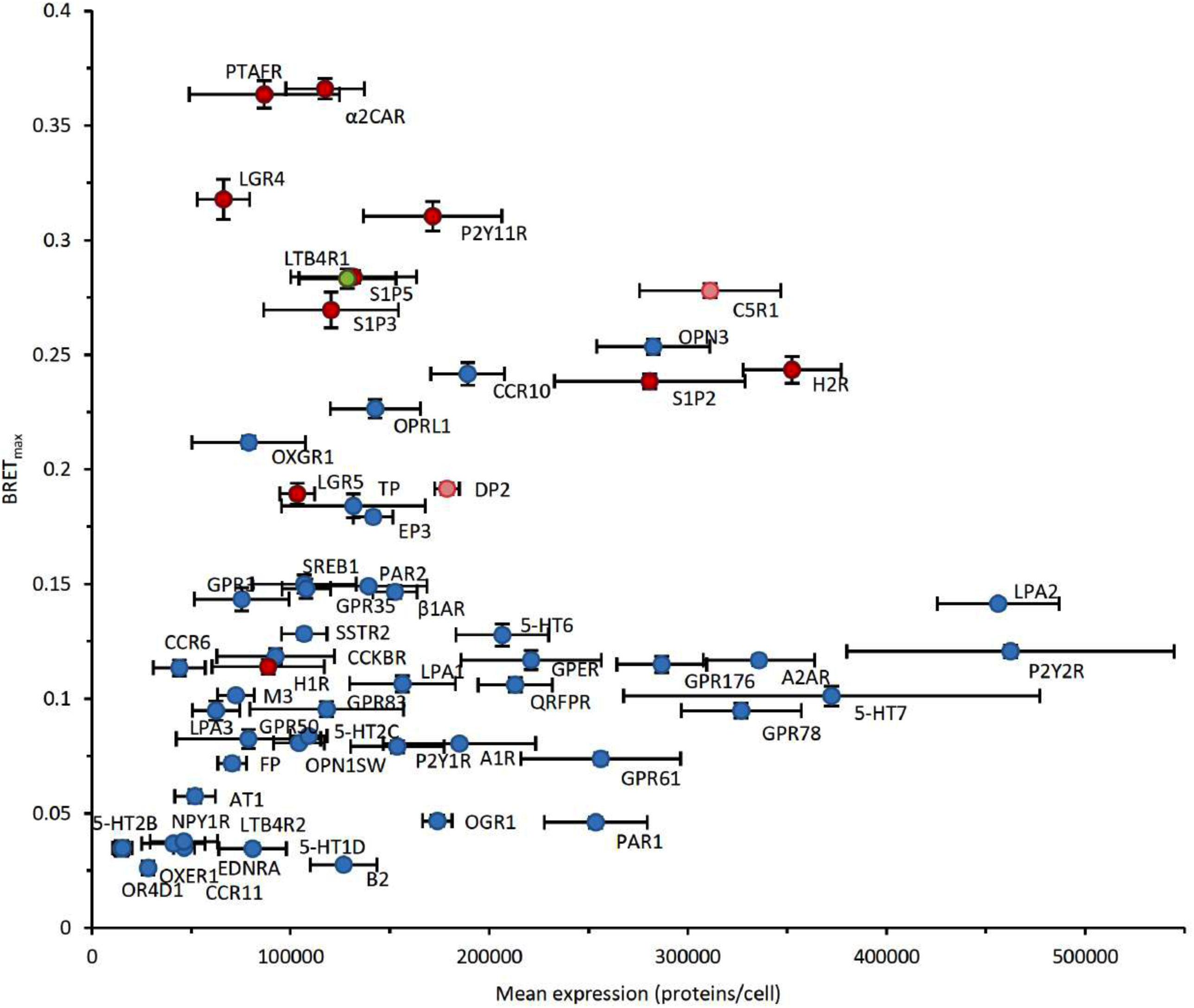
Relationship between BRET_max_ and expression level for HEK 293T *Rhodopsin*-family GPCRs. BRET_max_ values for monomers are generally lower than those of dimers at similar expression levels. Monomeric receptors are shown in blue; dimers in red. The ambiguous cases C5R1 and DP2 are shown in pink. LTB4R1 is shown in green. Protein names are located as close to their respective points as possible. Error bars indicate SEM. of each parameter. CXCR4 is not shown for clarity. Absolute values are given in Dataset S1.

**Figure S9.**
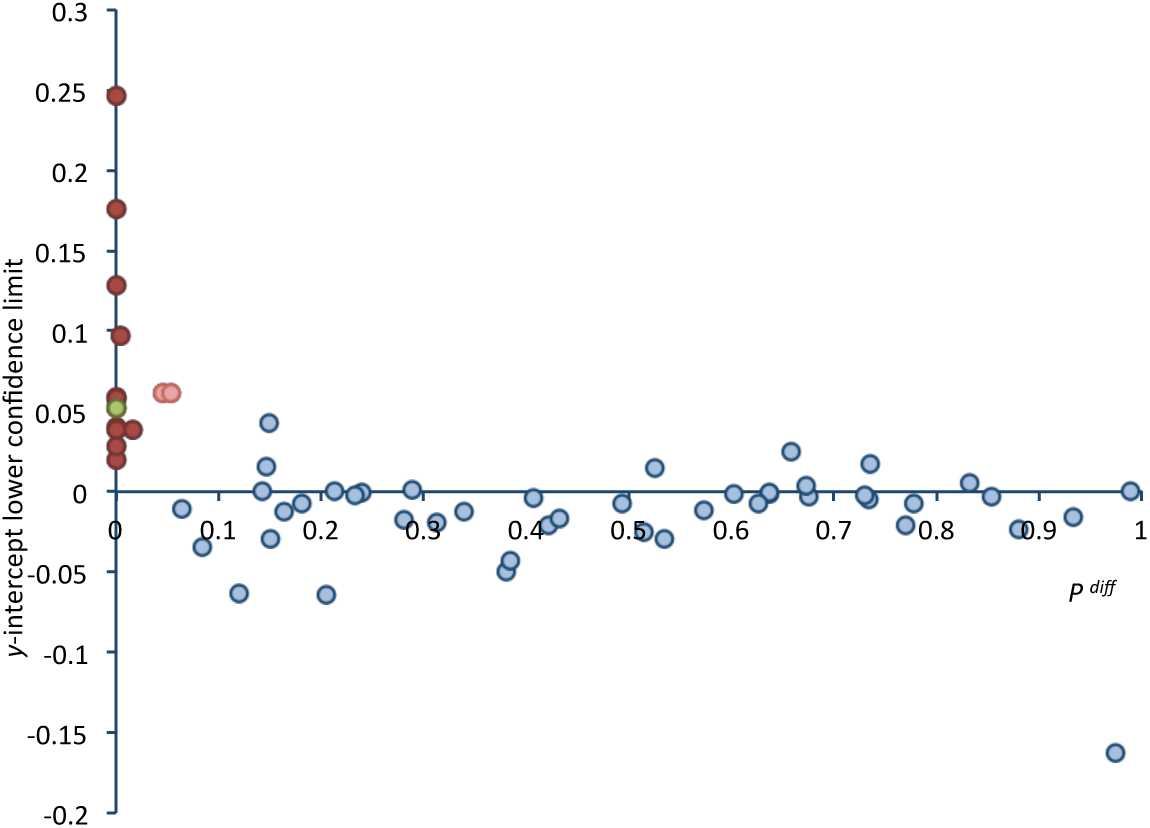
*y*-intercept values obtained using type-3 BRET in the absence of competitor typically correlated with apparent stoichiometry. Receptors concluded to be monomers are shown in blue, dimers in red. C5R1 and DP2 are shown in pink, LTB4R1 in green. All apparent *Rhodopsin-family* dimers yielded a *y*-intercept value with a lower 95% confidence limit that is above zero, as did the ambiguous cases C5R1, DP2, and LTB4R1. The majority of apparently monomeric receptors had y-intercept values that are not significantly non-zero, although a small number have lower 95% confidence limits greater than zero.

**Figure S10.**
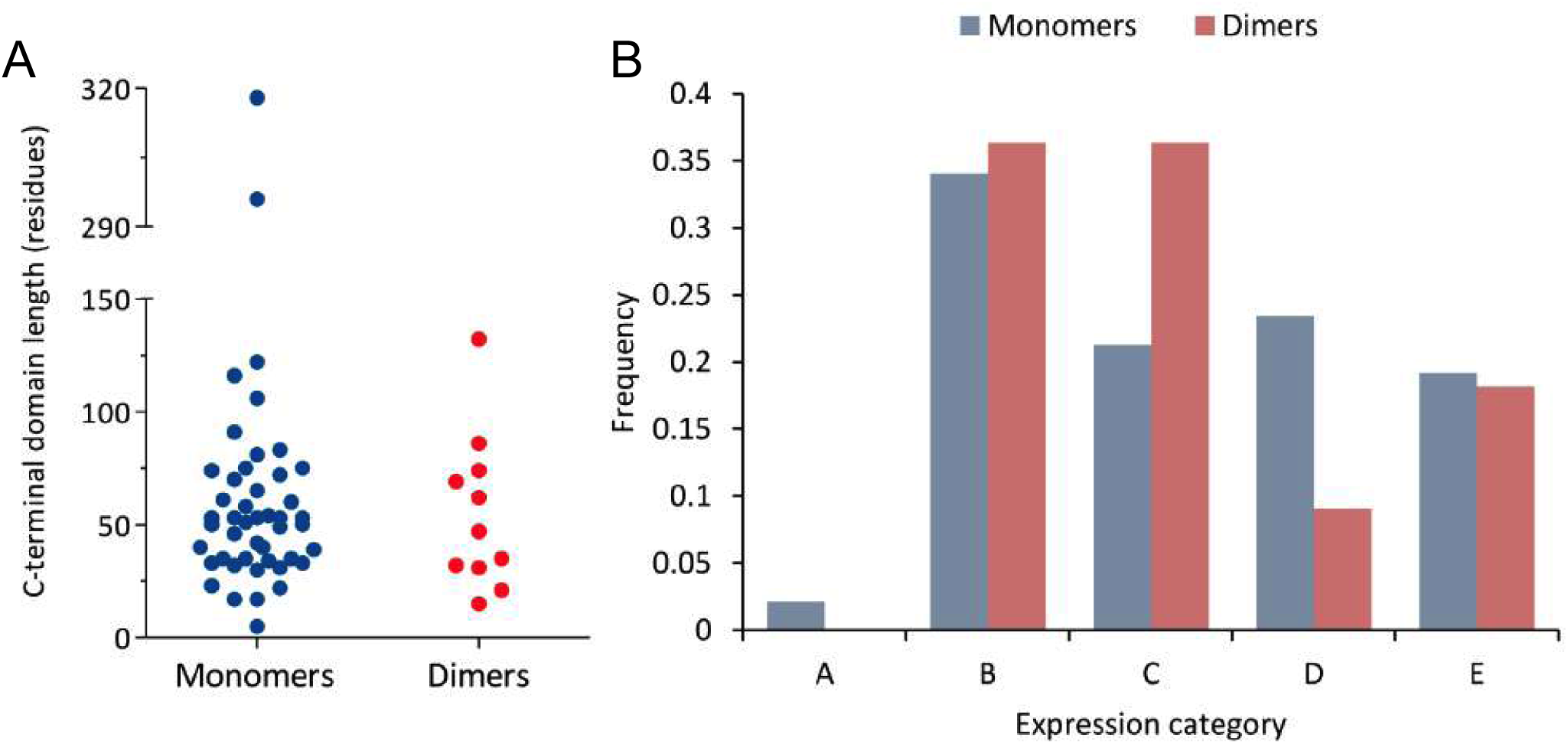
There is no clear difference in C-terminal topology or trafficking behavior between the *Rhodopsin*-family monomer and dimer populations. (A) Apparent monomers and dimers had comparable lengths of their C-terminal domains. This suggests that the observed monomers were not the mis-assignment of dimers with large C-terminal domains that preclude efficient energy transfer. (B) Apparent monomers and dimers demonstrated similar expression profiles. This indicates that apparent dimers were not artefacts of intracellular retention. C5R1, DP_2_, and LTB4R1 are not included. Expression categories correspond to those detailed in Table S1.

**Figure S11.**
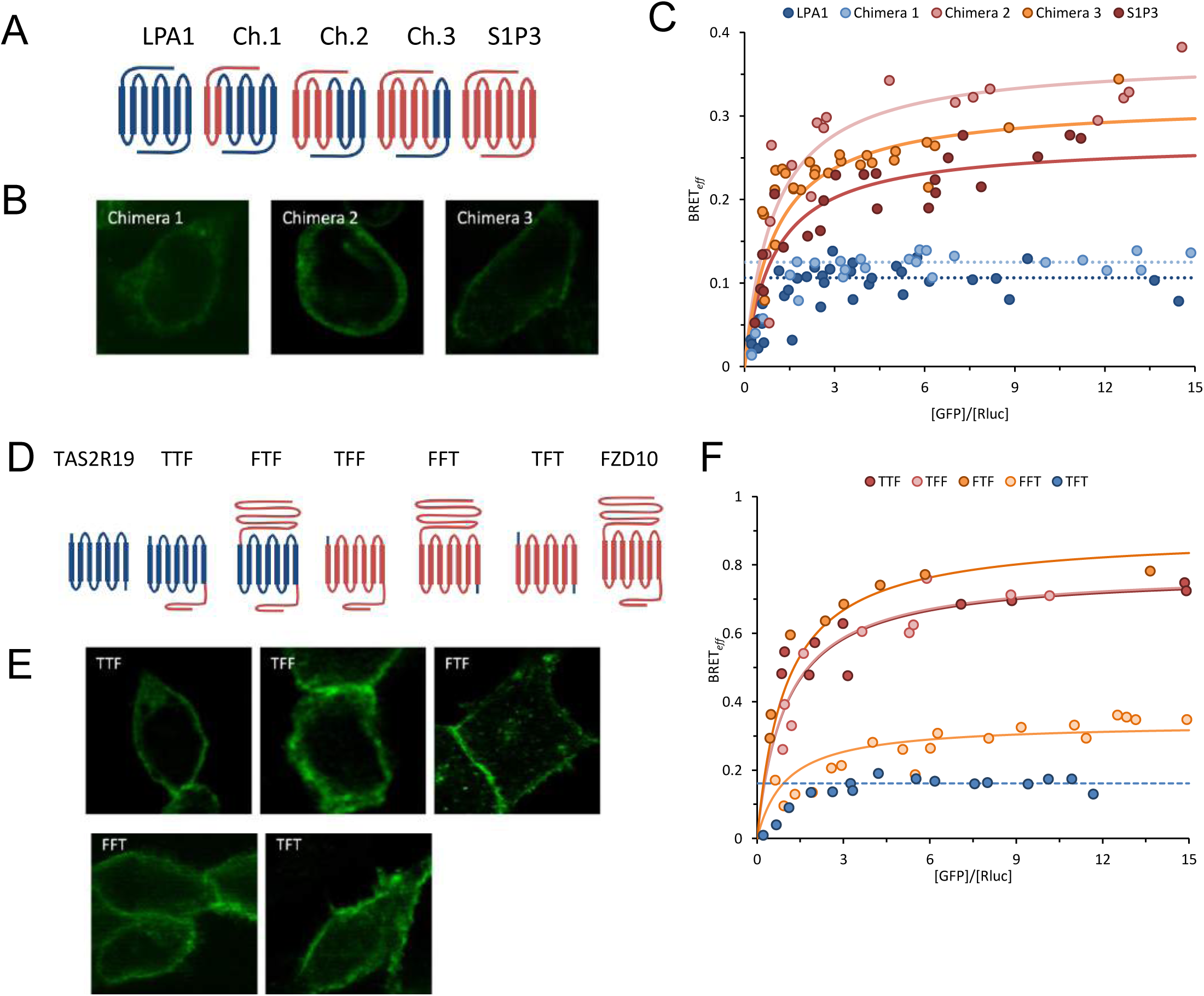
Analysis of chimeric receptors reveals different mechanisms of dimerization between *Rhodopsin*- and *Frizzled*-family receptors. (**A**) Schematic representations of the LPA1 (blue) and S1P3 (red) composition of each construct. (**B**) Representative confocal microscopy images of HEK 293T cells expressing the three LPA1-S1P3 chimeras from the pGFP^2^ vector. All three constructs exhibit increased intracellular retention compared to the parent genes, but not aggregation and so all were suitable for use in BRET. (**C**) Type-1 BRET analysis of three LPA1-S1P3 chimeras as well as the parent receptors for comparison. Chimera 1 exhibits an independence of BRET_*eff*_ upon [GFP]/[Rluc], indicative of a monomer. Fits of LPA1 and chimera 1 data to a constant model are shown as broken lines. Chimeras 2 and 3 demonstrate hyperbolic dependences of BRET_*eff*_ upon [GFP]/[Rluc] that fit well to a dimer model (solid lines). This indicates the S1P3 dimerization is dependent on motifs between EL1 and TM4. (**D**) Schematic representations of chimeras of TAS2R19 (blue) and FZD10 (red). Chimeras were given a three-letter designation based on their composition, in which F and T denote FZD10 and TAS2R19 components, and the first, second, and third letters indicate the origin of the N-terminal domain, TM region, and C-terminal domain, respectively. Of the six possible combinations, only FTT failed to express sufficiently for BRET analysis. (**E**) Representative confocal microscopy images of HEK 293T cells expressing the 5 successfully expressed TAS2R19-FZD10 chimeras from the pGFP^2^ vector. Aggregation was not apparent in any case and so all were suitable for use in BRET. (**F**) Type-1 BRET analysis of TAS2R19-FZD10 chimeras indicates a role in dimerization of both the FZD10 N- and C-terminal domains. All chimeras containing either the FZD10 N- or C-terminal domains (TTF, TFF, FTF, and FFT) exhibited BRET_*eff*_ dependence upon [GFP]/[Rluc] in the manner predicted for a dimer. Replacement of the N- and C-terminal domains of FZD10 with those of TAS2R19 (chimera TFT) caused BRET_*eff*_ to become independent of [GFP]/[Rluc], which indicates monomeric behavior. This suggests that the FZD10 TM region does not possess any inherent dimerization ability, in contrast to S1P3.

**Table S1.** Qualitative categories of GPCR expression based on cellular localization and aggregation.

| Category | Definition |
| --- | --- |
| A | Protein entirely in plasma membrane; almost no visible protein in internal membranes; no aggregation. |
| B | Protein almost entirely in plasma membrane; small amounts in internal membranes; no aggregation. |
| C | Majority of protein in plasma membrane; moderate amounts in internal membranes; no aggregation. |
| D | Some protein in plasma membrane; large amounts in internal membranes; no aggregation. |
| E | Some protein in plasma membrane; large amounts in internal membranes; small amounts of aggregation in some cells. |
| F | Some protein in plasma membrane; large amount in internal membranes; small amounts of aggregation in most cells. |
| G | Little or no protein in plasma membrane or internal membranes; large amounts of aggregation in most cells. |

**Table S2.** Transfection conditions, HaloTag density, and coincidence values for controls and GPCRs analyzed using SMCCCD in CHO K1 cells.

| Protein | HALO-tagged Construct (μg) | SNAP-tagged Construct (μg) | Post-transfection incubation (hours) | Cells imaged | Mean HaloTag spots/cell ± SD | Mean % coincidence ± SEM | *p*-value of difference from CD86 |
| --- | --- | --- | --- | --- | --- | --- | --- |
| CD86 | 0.975 | 0.175 | 20 | 10 | 311±138 | 9.7±1.5 | - |
| CD28 | 0.975 | 0.175 | 48 | 8 | 288±85 | 28.2±3.7 | 0.0001 |
| LPA1 | 0.975 | 0.08 | 24 | 9 | 312±194 | 13.8±2.6 | 0.173 |
| S1P3 | 0.975 | 0.08 | 20 | 9 | 422±150 | 22.9±3.4 | 0.002 |
| β<sub>1</sub>AR | 0.975 | 0.08 | 15 | 8 | 299±88 | 11.7±1.4 | 0.366 |
| α<sub>2C</sub>AR | 0.975 | 0.08 | 20 | 15 | 385±114 | 14.8±1.3 | 0.017 |

**Table S3.**
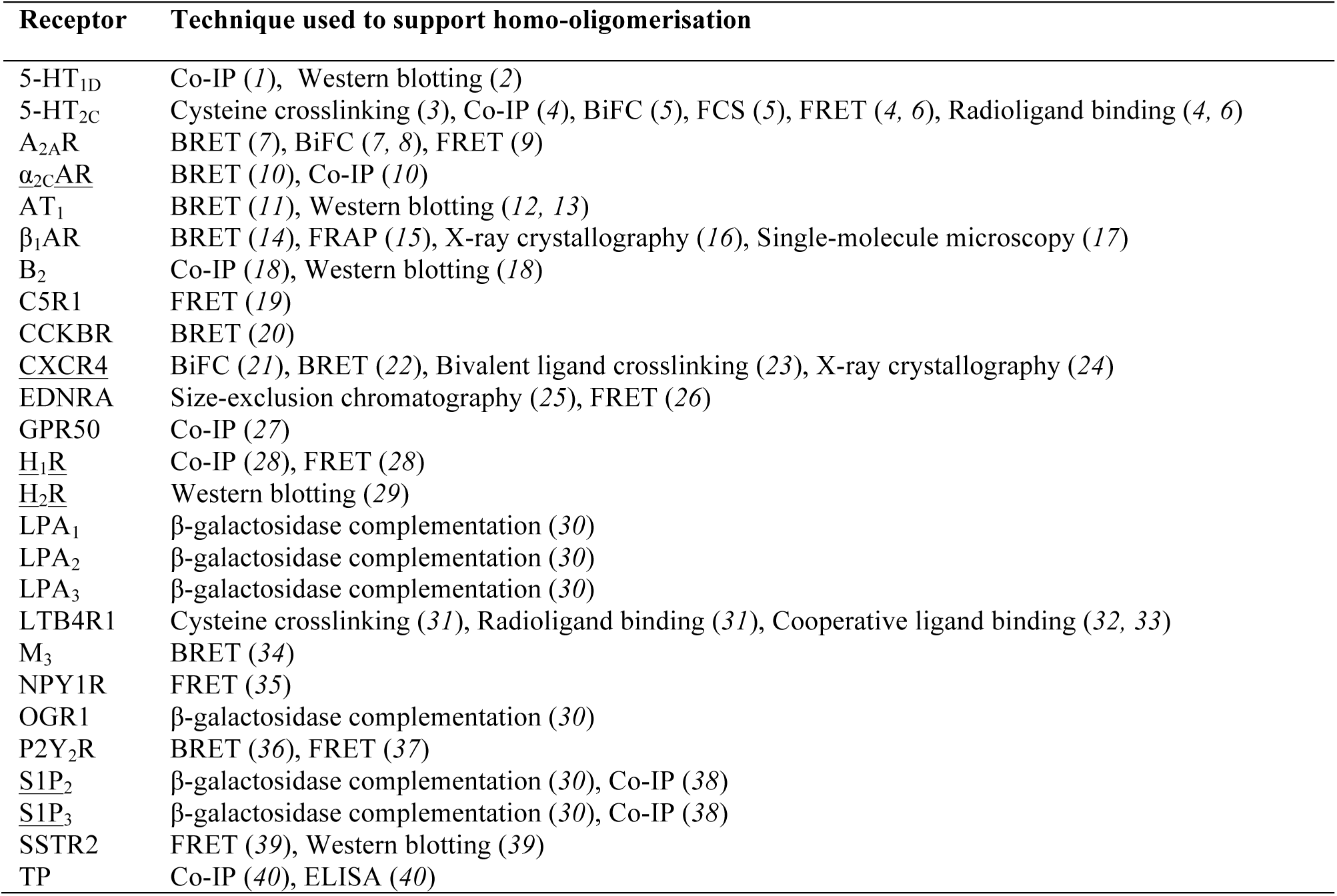
Previously published assertions of homo-oligomerisation by *Rhodopsin*-family GPCRs investigated in this study. 26 of the 60 receptors investigated have previously been reported as homodimers in studies using a range of techniques. GPCRs found to be dimeric using type-1 and -3 BRET assays are underlined. Only reports of homo-oligomerisation are included in this table; studies of heteromeric interactions are not shown.

**Dataset S1.**
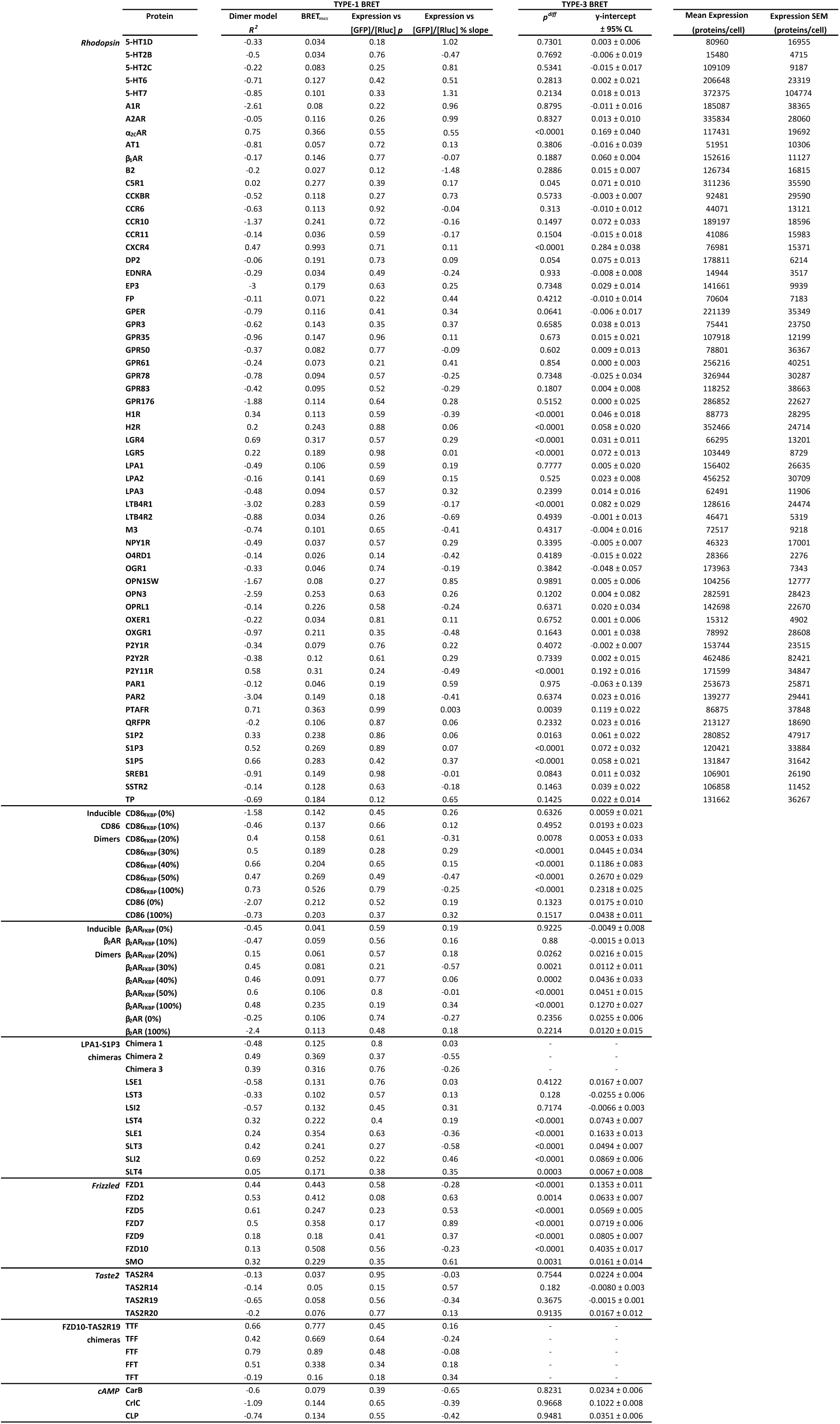
Raw outcomes for type-1 and -3 BRET assays and quantified expression levels for all investigated receptors.

**Dataset S2.**
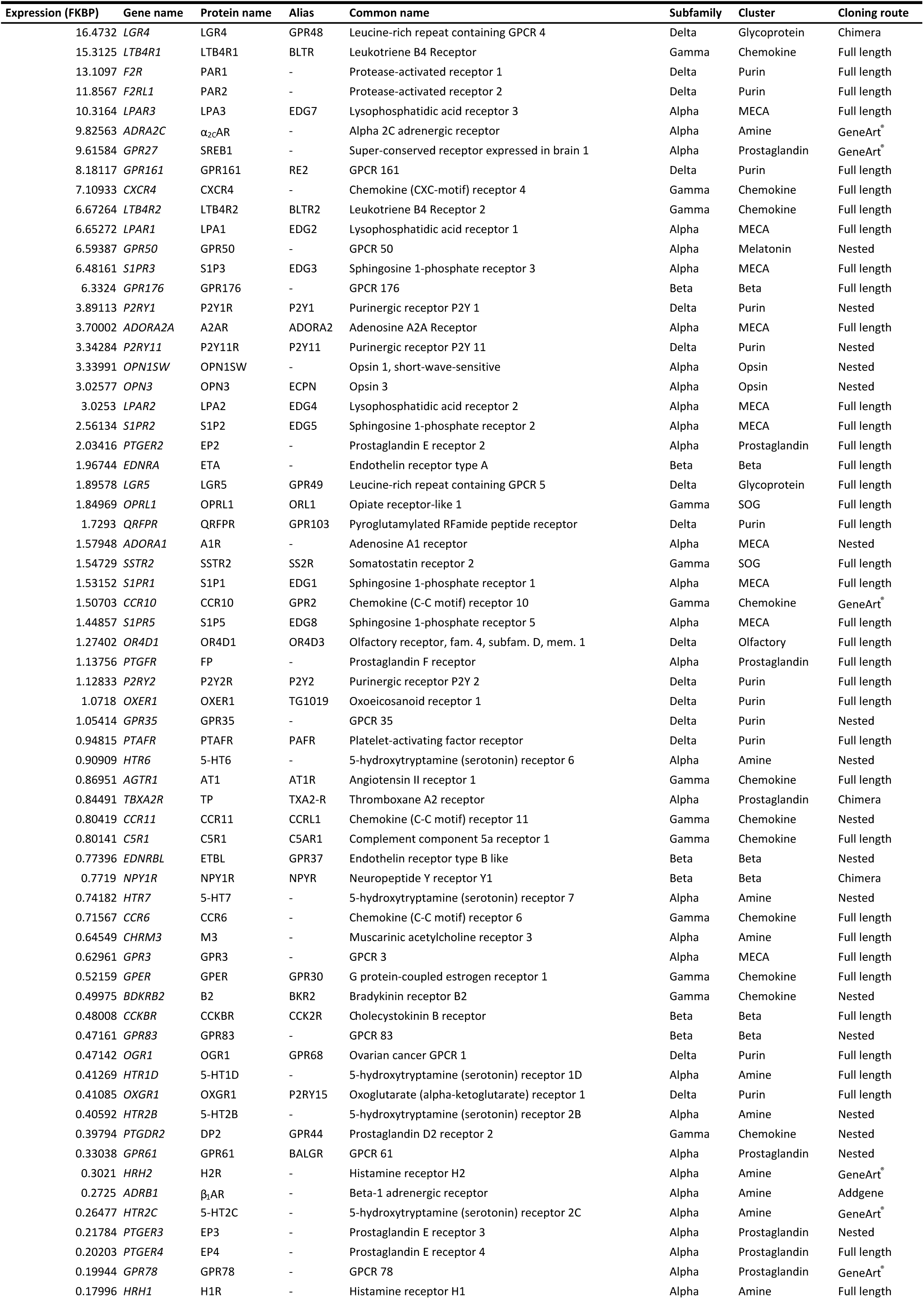
Full list of *Rhodopsin*-family GPCRs expressed in HEK 293T cells and the cloning strategy used in each case.

## Technical Note

### The type-1 BRET assay

In type-1 BRET, total protein concentration is kept constant and the ratio of GFP- to Rluc-tagged proteins is varied. For monomers, there is no dependency of BRET_*eff*_ on acceptor:donor ratio above a certain threshold (*1*) because the acceptor:donor ratio reaches a point at which the acceptors are in large excess over donors and so donors no longer compete for acceptors (Fig. S1). Thus, altering the ratio by removing donors and replacing them with acceptors has essentially no effect on the average distance between each remaining donor and its nearest acceptor, meaning that BRET_*eff*_ remains constant. In marked contrast, BRET_*eff*_ for dimers shows a strong dependence on acceptor:donor ratio because, as the ratio increases, fewer unproductive donor-donor pairs form and each donor is more likely to be paired with an acceptor, increasing BRET_*eff*_. This dependence can be expressed as a hyperbolic relationship between BRET_*eff*_ and acceptor:donor ratio (*f*) that is defined by the stoichiometry (*n*) of the oligomer (Equation 1). The derivation of this relationship has been published previously (*2*). The outcome of the type-1 assay is determined by the goodness-of-fit of the collected data to the dimer model and whether this is more accurate than a fit to the flat model predicted of monomers (Fig. S2).

To allow type-1 assay data to be reliably interpreted, total receptor density must not vary across different acceptor:donor ratios. Any systematic increase in density as acceptor:donor ratio increases may lead to a false dimer signature, whereas a decrease may lead to a false monomer signature. It is therefore important to calculate total receptor density at all acceptor:donor ratios, which can be done using the acceptor:donor ratio and absolute luminescence or fluorescence values (*i.e.* total expression = RLU + (RLU x [GFP]/[Rluc]); RLU = relative luminescence units).

Another important consideration in the use of the type-1 assay is that it does not discriminate between receptors in different cellular compartments or membranes. Tagged protein that is retained intracellularly does not interfere with the interpretation of type-1 assay data so long as it is not aggregated, since aggregation causes an effective oligomer signal that will mask any genuinely dimeric behavior and lead to a false monomer. Screening candidate proteins for aggregation is therefore a key precursor to the performance of BRET experiments.

### The type-3 BRET assay

The type-3 BRET approach is a modified version of the BRET competition assay. In such assays BRET_*eff*_ is measured in the presence and absence of untagged ‘competitor’ proteins; BRET between monomeric proteins should be unaffected, whereas BRET-productive dimers are disrupted by the competitors and thus BRET_*eff*_ is reduced (Fig. S1). The type-3 assay involves performing such measurements at varying receptor densities but a constant acceptor:donor ratio in order to remove the potential for artifactual decreases in BRET_*eff*_ arising from reduced expression of tagged proteins in response to competitor addition. The outcome of the assay is determined by fitting data collected in the presence and absence of competitor to a two-tailed *t* test of least-squares linear regression; a significant difference indicates dimerization or oligomerization (Fig. S3).

Importantly, the type-3 assay is not interpreted in the same manner as another assay, type-2 BRET (2), which is essentially analogous to a type-3 experiment in the absence of competitor. In a type-2 assay, stoichiometry is inferred from the *y*-intercept of a linear regression of the data as monomers regress to an intercept of zero whereas dimers have a non-zero intercept since BRET_*eff*_ within dimers is not density- dependent. In order to achieve a precise *y*-intercept, data are collected at the lowest possible expression levels that allow signal detection, which also ensures the BRET_*eff*_ expression relationship is linear. This makes the type-2 assay less suitable for the examination of receptors in cells with native expression of the protein of interest. The type-3 assay is performed at much higher receptor densities and thus is not vulnerable to contributions from native proteins. However, this also renders the y-intercept as a less reliable metric for interpreting stoichiometry because the slope of BRET_*eff*_ *vs.* expression can change at higher protein densities and so artefactually increase the *y*- intercept estimated by linear regression.

Like the type-1 assay, type-3 experiments are sensitive to protein aggregation, although in this case it will yield a false dimer outcome.

### Limitations of the type-1 and -3 BRET assays

Type-1 BRET relies on statistical comparisons with model behavior predicted for different classes of stoichiometries, whereas the type-3 assay simply measures the change in BRET_*eff*_-expression relationship upon the introduction of competitor molecules. As such, type-1 BRET is very unlikely to generate false-dimer artifacts due to the highly diagnostic nature of data distribution for dimers, but is potentially at risk of providing false-monomer results in cases of higher-order oligomerization or weak dimerization (Fig. S1). Conversely, the type-3 BRET assay is highly unlikely to give false-monomer results, as this would require a competitor-induced increase in tagged protein expression that precisely compensates for the drop in BRET_*eff*_ caused by competition. It is possible, however, that the type-3 approach could produce false dimers in certain cases where addition of competitor proteins causes a relaxation in the localization of tagged proteins within the membrane, thereby reducing their effective concentration and, by extension, BRET_*eff*_ (Fig. S1). Thus, complementary use of both type-1 and -3 BRET greatly increases confidence in assay outcomes, as any agreement between the two is extremely unlikely to be due to technical artifacts.

Although the combination of type-1 and -3 assays allows confident assignment of stoichiometry to transfected receptors, the need for receptor overexpression is an important consideration in the interpretation of BRET-derived data. Due to the laws of mass action, insignificant dimerization at native expression levels may be greatly stabilized as receptor density increases in transfected cells. It is always possible, therefore, that dimers observed under these conditions do not reflect their *in vivo* behavior. Proteins reported as monomers, however, are unlikely to represent a deviation from their native stoichiometry except in cases where association is dependent on a limited pool of some additional interaction partner. In the case of GPCRs, dimerization has typically been regarded as a fundamental chemical property of the receptors that does not require stabilization by accessory proteins.

The interpretation of BRET-derived data is also limited to a broadly binary ‘monomeric’ or ‘dimeric’ (>10-20% dimerization) delineation, and more nuanced interpretations are usually not possible. In the type-1 assay, data are compared between models of absolute monomeric and dimeric behavior. Fitting data to models of partial dimerization is in principle possible, however to be done with confidence it would require greater experimental precision than is currently feasible. The type-1 assay is also unable to distinguish between monomers and higher-order oligomers, which also deviate significantly from the distribution expected of dimers. Moreover, type-1 assay BRET_max_ cannot normally be directly interpreted as a measure of the extent of dimerization as it is determined by a wide range of variables, including receptor density, clustering, and geometry within the dimer. An exception to this is in cases of very high BRET_max_ (*i.e.* >0.9) as such levels of BRET can only feasibly be explained by robust association. In the type-3 assay, the magnitude of any reduction in BRET_*eff*_ in the presence of competitor does not necessarily correlate with strength of association since this is dependent on the extent to which dimers containing competitor receptors are able to undergo non-specific BRET with one another in the membrane.

### Single-molecule cross-color coincidence detection (SMCCCD)

SMCCCD requires the co-expression of the receptor of interest tagged separately with two enzymatic tags, SNAP-tag and HaloTag, which allow its simultaneous visualization in two colors (Fig. S1). In order to resolve single receptors, total internal reflection fluorescence microscopy is used, wherein only those fluorophores within approximately 100-200nm of the glass coverslip on which the sample is mounted are illuminated and so the background-signal contrast is sufficient to detect single fluorophores. Single-molecule resolution is also contingent on a low density of receptors, as high densities cause individual signals to become merged into an ensemble image. For this reason, expression of tagged receptors is performed under the control of a restricted promoter, which ensures low expression levels. This is in stark contrast to the BRET assays, which require overexpression. As a result, SMCCCD must be performed in cells that do not possess the native form of the receptor, which is not an important consideration in the type-1 and -3 BRET assays.

In the present SMCCCD experiments, receptors are expressed in CHO-K1 cells, which are then labeled with SNAP-tag and HaloTag ligands prior to fixation and imaging. Fixed cells are used because stoichiometry is inferred by comparison to known controls, and using live cells introduces the additional variable of dissimilar diffusive behavior between samples and controls (for a more detailed discussion of the requirement for fixation see Latty et al. (*3*)). Samples are imaged for 400 frames of 35 ms in order to allow elimination of the small fraction of still mobile receptors. Individual fluorescent puncta (corresponding to a single-diffraction limited spot) are then identified and classed as ‘coincident’ or ‘non-coincident’ based on their proximity to puncta in the opposing color. HaloTag-labeled receptors that remain within 300nm of one or more SNAP-tag-labeled receptors for 10 or more frames are deemed coincident (the selection of these values has been discussed previously (*3*)).

Unlike the BRET assays, SMCCCD data cannot be directly interpreted and instead relies on comparison to controls of known stoichiometry. This has the added effect of allowing more confident speculation of the extent of any dimerization by reference to the absolute coincidence values of samples and controls, in contrast to the largely binary outcome of the BRET approaches. To allow appropriate comparison between samples and controls protein density must be kept constant, as any change will result in differing levels of non-specific coincidence. This is achieved by screening for transfection conditions that lead to comparable densities and ratios of the SNAP-tag- and HaloTag-labeled forms of all proteins imaged. The efficiency of receptor labeling has been quantified previously (*3*), however in principle the outcome of SMCCCD is independent of labeling efficiency.

### Limitations of SMCCCD

A limitation of SMCCCD is that resolution is constrained by the Abbe diffraction limit. Fluorophores separated by a distance smaller than this limit cannot be independently resolved due to diffraction, and as a result they appear as single objects. The size of the diffraction limit (d) is determined by the wavelength emitted (λ) as well as the numerical aperture (NA) of the instrument according to the relationship d = λ /2NA, typically ~300 nm. This is ~6 times larger than the hydrodynamic diameter of most GPCRs and over 20 times larger than the effective resolution of most RET approaches, which is typically 5-10nm. It is therefore inherently impossible to distinguish between genuine dimerization and non-specific colocalization of receptors on a scale below the diffraction limit (Fig. S1). This possibility is minimized through the use of low receptor densities, however the most circumspect strategy is to compare SMCCCD-derived conclusions with those obtained using BRET assays, since their small effective resolution makes misidentification of stable non-specific colocalization unfeasible.

Comparison of key factors in the interpretation of BRET and SMCCCD experiments.

|  | Type-1 BRET | Type-3 BRET | SMCCCD |
| --- | --- | --- | --- |
| **Effective resolution** | 10nm | 10nm | 300nm |
| **Receptors/cell** | 100,000-500,000 | 100,000-500,000 | 1,000-5,000 |
| **Cell type** | Native | Native | Non-native |
| **Cell condition** | Live | Live | Fixed |
| **Distinguish monomers vs. dimers?** | Yes | Yes | Yes |
| **Distinguish monomers vs. oligomers?** | No | Yes | Yes |
| **Distinguish oligomers vs. dimers?** | Yes | No | No |
| **Distinguish dimers vs. colocalization?** | Yes | Yes | No |
| **Affected by intracellular aggregation?** | Yes | Yes | No |
| **Potential for false dimers?** | No | Yes | Yes |
| **Potential for false monomers?** | Yes | No | No |
| **Conclusion** | Binary | Binary | Continuous |

1. B. K. K. Fung, L. Stryer, Surface density determination in membranes by fluorescence energy transfer. *Biochemistry* **17**, 5241-5248 (1978).
2. J. R. James, M. I. Oliveira, A. M. Carmo, A. Iaboni, S. J. Davis, A rigorous experimental framework for detecting protein oligomerization using bioluminescence resonance energy transfer. *Nat. Methods* **3**, 1001-1006 (2006).
3. S. L. Latty *et al.*, Referenced single-molecule measurements differentiate between GPCR oligomerization states. *Biophys. J.* **109**, 1798-1806 (2015).

